# Dynamic ensembles of SARS-CoV-2 N-protein reveal head-to-head coiled-coil-driven oligomerization and phase separation

**DOI:** 10.1101/2024.12.02.626213

**Authors:** Guillem Hernandez, Maria L. Martins, Nuno P. Fernandes, Tiago Veloso, João Lopes, Tiago Gomes, Tiago N. Cordeiro

## Abstract

The SARS-CoV-2 nucleocapsid (N) protein is essential for the viral lifecycle, facilitating RNA packaging, replication, and host-cell interactions. Its ability to self-assemble and undergo liquid-liquid phase separation (LLPS) is critical for these functions but remains poorly understood. Using an integrated approach combining small-angle X-ray scattering (SAXS), nuclear magnetic resonance (NMR) spectroscopy, computational modeling, and biophysical assays, we uncover key mechanisms underpinning N-protein’s dynamic self-assembly. We show that the N-protein’s interdomain linker (IDL) contains a conserved coiled-coil (CC) motif that drives transient interactions between protein subunits, enabling the formation of progressively larger complexes at higher concentrations. SAXS analysis and ensemble modeling reveal that the IDL exists in a concentration-dependent equilibrium between monomeric, dimeric, and trimeric states. The CC motif facilitates parallel, head-to-head oligomerization of N-protein dimers, transitioning between compact (closed) and extended (open) configurations depending on the interaction network within the IDL. This linker-driven assembly modulates LLPS, impacting the size, stability, and dynamics of biomolecular condensates. Here, we present the most comprehensive conformational landscape analysis of the N-protein to date, providing a detailed model of its self-assembly and LLPS. Our findings highlight how the structural plasticity of the IDL and CC-mediated interactions are pivotal to its roles in the SARS-CoV-2 lifecycle.

## INTRODUCTION

Coronaviruses (CoVs) are a diverse family of viruses that infect mammals and birds, with seven strains known to infect humans. While some, such as HCoV-OC43 and HCoV-HKU1, cause mild respiratory tract infections, others—like MERS-CoV, SARS-CoV-1, and the global pandemic SARS-CoV-2—can lead to severe and potentially fatal diseases (1). These outbreaks highlight the critical need to unravel the molecular mechanisms underlying CoV infection and pathogenesis.

CoVs are single-stranded RNA viruses with a genome ranging from 27 to 34 kb (2). Packed inside the virion by the nucleocapsid protein (N-protein), their genome encodes the virus’s replication machinery and essential viral proteins. The prominent SARS-CoV-2 has a 30-kb-long genome encoding about 29 proteins. Among these proteins, the N-protein stands out as one of the most highly expressed within host-infected cells (3), serving as a relevant marker of infection and a key focus for SARS-CoV-2 vaccine development (4, 5). However, its high immunogenicity causes an imbalance between immunity and immunopathogenesis, and further studies are needed to harness its therapeutic potential (6).

In addition to encapsulating the sizable viral genome RNA (gRNA) inside the 80-90nm virion (7), N-protein performs several other vital functions essential for the SARS-CoV-2 lifecycle. It protects the gRNA in the host cell environment (8), participates in its transcription regulation (9, 10), influences host-cell responses (1, 11, 12), and acts as a vital cofactor for virus replication machinery and assembly (7). Emerging evidence shows that the N protein performs these functions via self-assembly and liquid-liquid phase separation (LLPS) (13–17).

Upon viral entry, SARS-CoV-2 generates double-membrane vesicles (DMVs) from the endoplasmic reticulum into the cytosol (18), where RNA replication occurs. The RNA is then enveloped by N-protein, forming viral ribonucleoprotein (vRNP) complexes (19), which are then incorporated into new virions. The SARS-CoV-2 N-protein self-assembles and undergoes LLPS with RNA (17) for assembling vRNP complexes and potentially facilitating viral assembly (17). LLPS, a phenomenon in which biomolecules spontaneously form liquid-like droplets or gels within cellular environments (20), plays a crucial role in modulating the innate antiviral immune response. It can achieve this by dampening IFN-β signaling (21) or triggering NF-κB signaling (22). Therefore, targeting the LLPS of the N-protein could be a potential therapeutic strategy against COVID-19.

The N-protein’s ability to self-assemble and mediate LLPS is facilitated by its dynamic modular architecture, which includes several intrinsically disordered regions (IDRs). Specifically, the N-protein comprises two globular domains, N-terminal domain (NTD) and C-terminal domain (CTD), connected by a conserved interdomain disordered linker (IDL), with additional IDRs located at the N- and C-termini (13, 23). The NTD is responsible for RNA binding, while the CTD facilitates protein dimerization through a domain swap mechanism, which also involves RNA binding. These structural characteristics contribute to the protein’s structural flexibility and ability to bind to multiple targets, enabling it to form condensates.

The IDRs of the N-protein are particularly important for LLPS. The central IDL is critical for robust RNA-dependent N-protein phase separation (17), and the phosphorylation of its Arginine/Serine(SR)-enriched region modulates the biophysical properties of the N:RNA condensates (17, 24), thereby influencing their functions. For instance, gel-like condensates promote nucleocapsid assembly, whereas liquid-like condensates facilitate viral genome processing (24). The SR-rich motif is adjacent to a Leucine/Glutamine (LQ) region (residues 210-246) (17), harbouring a Leucine-rich helix (^219^LALLLLDRLNQLE^231^) (LR) that binds its viral partner Nsp3a (23) and participates in oligomerization (25).

The N protein’s oligomerization and LLPS are crucial in the virus’s lifecycle. Understanding the principles underlying N-protein high-order structures and condensates is instrumental in unveiling SARS-CoV-2 biology and offering opportunities for antiviral strategies (26) that target the LLPS of the SARS-CoV-2 N protein (21). Despite significant progress and active research, the complex structural aspects of N-protein high-order assembly and phase separation remain to be fully elucidated. Specifically, the dynamic interplay between its distinct domains, the conformational diversity within its high-order structures, and the molecular determinants driving self-assembly and LLPS require further investigation.

This study delves into the structural dynamics of the N-protein’s IDL and its potential implications for viral function based on macromolecular assembly. Utilising nuclear magnetic resonance (NMR) spectroscopy, small-angle X-ray scattering (SAXS), molecular dynamics (MD) simulations, and ensemble molecular modelling, we aimed to provide comprehensive insights into the structural and functional attributes of the N-protein. Our integrative approach revealed that the N-protein’s IDL exists in a dynamic equilibrium between monomeric, dimeric, and trimeric states. This equilibrium contributes to the oligomerization of the full-length N-protein, which forms flexible tetramers via parallel coiled-coil interactions facilitated by the highly conserved leucine-rich helix within the IDL, harbouring a trimerization motif (RhxxhE) (27).

We propose that both “head-to-head” interactions at the IDL coiled-coil site and “tail-to-tail” interactions involving the CTDs contribute to forming a chain of linked N-protein molecules, leading to the assembly of high-order oligomers. Notably, we found that mutations abolishing the coiled-coil site (CC site) disrupt the formation of these oligomers and impact protein-mediated LLPS, highlighting the importance of this structural motif in oligomerization and droplet formation. Furthermore, our observations indicate that the NTDs exhibit flexible tethering to the CC sites, extending away from the core of the oligomers. This structural arrangement provides insights into the mechanism underlying N-protein oligomer formation and suggests potential functional roles for these oligomers during the viral lifecycle.

By analyzing 26 SAXS datasets across multiple constructs and generating 87 optimized structural ensembles spanning various oligomeric states, we present what we believe is the most comprehensive ensemble analysis of the N-protein to date. This extensive study enabled us to resolve its structural heterogeneity in solution, providing unique insights into its self-assembly mechanisms and phase-separation behavior.

## MATERIAL AND METHODS

### Intrinsic disorder, conservation, and structure prediction

Sequences of N-protein variants (ca. 212000) were retrieved from the NCBI SARS-CoV-2 Data Hub (www.ncbi.nlm.nih.gov/labs/virus), for taxid:2697049. Disorder propensities and the per-residue predicted local difference test (pLDDT) were computed with Metapredict (28) for every sequence, and the per-residue Shannon entropy was retrieved from the Nextstrain website (www.nextstrain.org) (29). Sequence Logos were calculated with WebLogo using default settings (30). We fed it with a multiple sequence alignment obtained with Consurf (31) using the N-protein from SARS-CoV-2 (UniProt: P0DTC9) as seed (32). We also generated a sequence Logo for variants of interest (VOI) obtained from a dedicated database (https://covariants.org/) (33). We predicted the propensity for secondary structure of N-protein’s IDL using PSIPRED (34).

We used AlphaFold-2 (AF-2) (35) to predict the structure of the linker region 215-240 by feeding the sequence GDAALALLLLDRLNQLESKMSGKGQQ (N_215-240_). AF-2 predictions were performed using the ColabFold implementation (36) installed locally and MMseqs2 to create multiple sequence alignments (37). We did not use template structures in the predictions, iterated for up to 48 recycles, followed by energy refinement with AMBER using default settings. We used AF-2-multimer (38) to predict dimer (CC-Di) and trimer (CC-Tri) coiled-coil structures of N_215-240_, a refined version of AF-2 for complex prediction. The modelling confidence was assessed using the pLDDT metric and the inter-chain predicted alignment error (PAE), i.e., the uncertainty about the interface. pLDDT is closely related to the pre-existing metric lDDT-Cα (39) that measures the local accuracy of a prediction by determining the fraction of preserved local distances (higher is better). As a superposition-free method, lDDT is insensitive to relative domain orientation and correctly identifies segments in the full-length model deviating from the reference structure. pLDDT scales from 0-100. Values of pLDDT > 90 imply high accuracy. PAE is not an inter-residue distance map or a contact map but the expected distance error in Å. It indicates the expected positional error at residue x if the predicted and actual structures are aligned on residue y (using the C, N, and C atoms). PAEs are measured in Å and capped at 30 Å. PAE helps to assess the confidence in the relative position and orientation of the model parts (e.g., two domains or chains). For residues x and y in two chains, if the PAE values (x, y) are low, AF-2 predicts the chains to have well-defined relative positions and orientations (40). The knobs-into-holes (KIH) packing was analyzed with iSOCKET (41).

### Circular Dichroism

A synthetic peptide containing the predicted N-protein’s helical region (residues 215 to 240) with >95% purity on high-performance liquid chromatography (HPLC) was purchased from NZYTech to perform Far-UV CD spectroscopy in a J-815 spectrophotometer (Jasco) using a 1-mm optical pathlength cuvette for high performance (QS, Hellma). The N_215-240_ peptide single far-UV spectra were recorded by averaging five accumulations ranging from 198 to 280nm, using a 0.1nm data pitch and 2nm excitement bandwidth at a scan speed of 50nm/min under several concentrations (28.5, 57, 75, 100, 200 and 400µM) at 5°C using NAse-free MilliQ water as solvent. The thermal denaturation experiment was followed by monitoring temperature-dependent stepwise changes in spectral features ranging from 5 to 95 °C in 5 °C steps in the 28.5 ***μ***M sample. The CD data were normalized over concentration and peptide chain length and used as input for the DichroWeb web server employing SELCON3 (42, 43) with Set 4 and Set 7 (44, 45) as reference sets methods to predict secondary content.

### Molecular Dynamics simulations

We performed all-atom molecular dynamics (MD) simulations with GROMACS 5.1.1 (46) using the Amber99sb-ildn force field (47) with TIP3P water. MD simulations consisted of N_215-240_ in dimer parallel or antiparallel orientation and trimeric coiled-coil parallel configuration (CC-Tri) to provide an independent insight into the observed coiled-coil (CC) interactions predicted with AF-2. Using the same heptad register as in the AF-2 models, we created starting CC models for the interdomain helix with CCBuilder 2.0 (48) with acetylated and amidated N and C termini, respectively. The CC-models were then inserted into a dodecahedron box, with dimensions 63.0 × 63.0 × 44.5 Å for CC-Di and 66.0 × 66.0 × 46.6 Å for CC-Tri. Standard periodic boundary conditions applied in all directions. The ionic strength was 150 mM NaCl at neutral pH to assure charge neutrality. The system was minimised for a maximum of 50000 steps or until the force constant was less than 1000 kJ/mol·nm. Energy minimization used the steepest-descent algorithm. The Particle mesh Ewald method was used for the non-bonded interactions with the cutoff distance of 10 Å. Before the final production simulation, the system was equilibrated, during 10 ns, using the NPT ensemble, followed by the NVT ensemble. Finally, we ran the MDs in duplicate for 1.5 μs with a 2 fs integration step. Temperature coupling was done with the Nose–Hoover algorithm at 300 K. Pressure coupling was done with the Parrinello–Rahman algorithm at 1 bar.

From the trajectories, root-mean-square fluctuation (RMSF) and native contact analysis (Q) (49) were calculated using MDAnalysis (50). The figures were generated using Chimera (51). Contact maps were calculated using a 4.5 Å cutoff, with routines implemented in MDTraj (52).

### Protein expression and Purification

The genes encoding for full-length and truncated versions of N-protein (Uniprot: P0DTC9) were PCR-amplified from a vector with the cDNA of N-protein acquired from Addgene (pGBW-m4046785; Plasmid #145684). The CC null mutants in which L223, L227, and L230 were mutated to prolines (L3P) were amplified from a synthetic *E. coli* codon-optimized gene of the full-length protein L3P mutant purchased from GenScript, using the primers in **Supplementary Table 1**). The PCR products encoding N_176-246_, N_1-246_, N_1-365_, and N_1-419_ WT (residues 176-246, 1-246, 1-365 and 1-419, respectively; **Supplementary Table 2**) and L3P variants (CC-null) were inserted into the IPTG-inducible pHTP8 vector (NZYTech) fused to an N-terminal TrxA-His_6_ tag and a C-terminal Strep-tag. GFP-tagged N_1-419_ WT and N_1-419_L3P constructs were created using the pHTP9 vector (NZYtech) for fluorescence microscopy studies. All constructs encoded HRV-3C protease recognition sites to remove the TrxA-His_6_ or GFP tags.

For protein production, we transformed *E. coli* BL21 (DE3) pLysS competent cells with the pHTP8 or pHTP9-based constructs and grown them at 37°C to a mid-log phase (OD_600_∼0.8-1.0) in LB media containing kanamycin (50 μg/mL) and chloramphenicol (34 μg/mL). We induced protein expression at 20°C with 0.25 mM IPTG for 18-20h, then we pelleted the cells at 4°C and stored them at −20°C. For having uniformly isotopic-labelled protein, before the induction cells were collected and transferred to M9 media (33.7 mM Na_2_HPO_4_, 22 mM KH_2_PO_4_, 8.6 mM NaCl, 1 mM MgSO_4_, 0.3mM CaCl_2_, 1 mg/L biotin, 1 mg/L of thiamine and trace elements as 134 µM EDTA, 31 µM FeCl_3_·6·H_2_O, 6.2 µM ZnCl_2_, 760 nM CuCl_2_·2H_2_O, 430 nM CoCl_2_·2H_2_O, 1.6 µM H_3_BO_3_, 81 nM MnCl_2_·4H_2_O, 52 nM Na_2_MoO_4_·2H_2_O) with 34 µg/mL of chloramphenicol and 50µg/mL of kanamycin and supplemented with 1g/L of ^15^NH_4_Cl and 2 g/L of ^13^C-U-glucose as solely nitrogen and carbon source.

For pHTP8 constructs, frozen cell pellets were thawed and resuspended in lysis buffer (100 mM Tris·HCl, 500 mM NaCl, 6 M Urea, 20mM Imidazole, 5 mM MgCl_2_, 5 mM beta-mercaptoethanol, 10% glycerol, pH=8.0) supplemented with Benzonase, DNAse I, EDTA-free protease cocktail inhibitor (PIC) (Sigma), and lysozyme and then lysed in a French Press. The lysates were clarified by centrifugation for 45 min at 42000 rpm and 4°C. Soluble fractions were diluted to double their volume and loaded into a 5 mL Hitrap FF crude (Cytiva) affinity column pre-equilibrated in the lysis buffer. TrxA-His_6_-fused proteins were eluted using the elution buffer (100 mM Tris·HCl, 500 mM NaCl, 3 M Urea, 500 mM Imidazole, 2 mM DTT, pH=8.0) and analyzed by SDS-PAGE. The elution fractions containing the fused protein were dialyzed against the dialysis buffer (100 mM Tris·HCl, 500 mM NaCl, 1 mM EDTA, 1 mM DTT, pH=8.0) overnight at 4°C using a dialysis SnakeSkin^TM^ (Thermo Scientific) (3.5 KMWCO). The dialysis membrane was then transferred into a fresh dialysis buffer supplemented with HRV-3C protease (1:100 protease:target protein ratio). After overnight incubation at 4°C, we assessed the Trx-His_6_ tag cleavage by SDS-PAGE. The solution was then loaded into a 5mL StrepTactin™XT 4Flow™ high-capacity columns (IBA Lifesciences) equilibrated in ST binding buffer (100 mM Tris·HCl, 500 mM NaCl, pH=8.0) to separate the Strep-tagged protein and the cleaved Trx-His_6_-tag discarded in the flowthrough. We eluted the Strep-tagged protein with ST buffer supplemented with 50 mM of Biotin. Fractions containing pure protein were passed through a HisTrap FF crude column previously equilibrated with IMAC binding buffer (100 mM Tris·HCl pH=8.0, 500 mM NaCl) to trap traces of non-cleaved Trx-His_6_ protein. Then, pure fractions were pooled and purified using a size-exclusion Superdex 200 10/300 increase column (Cytivia) for N_1-365_ and N_1-419_ or a Superdex 75 10/30 Increase column (Cytivia) for N_1-246_, N_176-246_. We used SDS-PAGE to control the quality of the fractions corresponding to the protein peaks. Samples were concentrated and stored at −20°C with PIC (Sigma) to avoid degradation.

For the pHTP9 GFP-tagged constructs, frozen cell pellets were resuspended in lysis buffer (100 mM Tris·HCl, 500 mM NaCl, 5 mM beta-mercaptoethanol, DNAse, lysozyme, PIC (Sigma), 1 mM EDTA, pH=8.0) and lysed as described above for the pHTP8 variants. Supernatants were collected and applied on a StrepTactin™XT 4Flow™ high capacity 5mL column. After washing, GFP-tagged protein was eluted with biotin elution buffer (100 mM Tris·HCl, 500 mM NaCl, 5 mM beta-mercaptoethanol, 50 mM biotin, pH=8.0). The eluted fractions were diluted to 300mM NaCl, with the biotin elution buffer without NaCl, and subsequently passed through an HiTrapTM Heparin HP 1mL column (Cytiva) to avoid possible DNA contaminations. The proteins were eluted in heparin elution buffer (100 mM Tris·HCl, 1 M NaCl, 5 mM beta-mercaptoethanol, pH=8.0). Pooled fractions were further purified using a size-exclusion Superdex 200 10/300 increase (Cytiva) chromatography column in size-exclusion buffer (100 mM Tris·HCl, 500 mM NaCl, 5 mM beta-mercaptoethanol, 1 mM EDTA, pH=7.2) and frozen at −80°C until further use.

### NMR spectroscopy

We recorded all NMR experiments at 298 K on a Bruker Avance II+ spectrometer operating at 18.8T (800 MHz) equipped with a helium cold TXI-cryoprobe. Spectra were processed with TopSpin, MDDNMR (53) (available at NMRBox (54)), and NMRPipe (55) and analysed using CARA (56). To assign the backbone of IDL (N_176-246_), we used triple-resonance Best-TROSY-based 3D experiments (57) [HNCO, HN(CA)CO, HNCA, HN(CO)CA, HNCACB, HN(CO)CACB and (H)N(CACO)NH] all recorded with nonuniform sampling on the 42 μM U-[^15^N,^13^C]-labelled protein in the NMR buffer (50mM Potassium Phosphate pH=6.5, 250mM NaCl, 8% D2O, 20µM 3-trimethylsilyl-1-propanesufonic acid sodium salt (DSS)). Proton dimensions were directly corrected using the DSS signal (methyl group signal at 0.00ppm), while ^15^N and ^13^C were indirectly calibrated using the proton calibration. We computed the secondary structure propensities in terms of α-helix or β-sheet from the assigned chemical shifts using N-TALOS (58). Several 2D [^15^N-^1^HN]-BEST-TROSY were acquired on 30, 42, 85 and 182 μM samples of U-^15^N/^13^C-labelled N_176-246_ in the NMR buffer to monitor the effect of concentration. Moreover, T2 relaxation measurements were obtained using standard pulse sequences at 283K on the 30, 42 and 85 μM samples. We set the number of loops at 1, 2, 4, 8, 10, 12, 14, 16, 20, 24, 30, 40 and computed the relaxation rates by applying a one-phase decay function to the peak intensities. Relaxation rate errors were derived from the residuals’ covariance matrix calculated with the least squares method implemented in CcpNmr AnalysisAssign (59).

### Small angle X-ray scattering

We employed synchrotron SAXS to probe the overall size, conformational properties and oligomeric states of the different N-protein variants. SAXS data for N_176-246_, N_1-246_, N_1-365_ and N_1-419_ wild-type and L3P mutants were collected in B21 beamline of the Diamond Light Source (DLS, Didcot, UK) (60) and BM29 beamline of the European Synchrotron Radiation Facility (ESRF, Grenoble, France) (61), exploiting their unique in-line systems using a EigerX 4M (DLS) and Pilatus 1M (ESRF) detectors at a sample-detector distance of 3.7 m (DLS) and 2.8 m (ESRF) and at wavelengths of 0.954 nm (DLS) and 0.999 nm (ESRF) (I(s) vs s, where s = 4πsinθ/λ, and 2θ is the scattering angle). We injected 60 μL samples of growing concentrations in batch mode ranging from 0.9 to 12.0 mg/mL in 100mM Tris·HCl pH=8.0, 150mM NaCl, 1mM DTT, and 1mM EDTA, collecting 2-second frames at 10°C. We did not detect radiation damage. Data were integrated, buffer subtracted, and averaged to deliver SAXS profiles and further processed using PRIMUS from the ATSAS (62) package. From these subtracted data, the pairwise distance distribution functions, P(r), were obtained by indirect Fourier Transform with GNOM 5.0 available from ATSAS. We extracted the Rg values using the Guinier approximation in the range s < 1.3/Rg. Details are in **Supplementary Tables 3 and 4**.

### Structural ensembles

Ensemble models representing the monomeric, dimeric and trimer states of the N-protein interdomain linker (IDL, N_176-246_) were constructed using the Flexible-Meccano (FM) algorithm (63). The FM algorithm allowed us to generate random coil conformational ensembles for the disordered fragments flanking helical region (H) of the IDL. For these ensemble models, the conformation of H was fixed to a helix as defined by NMR constraints or to the predicted dimer and trimer models obtained from AF-2. This hybrid approach has been successfully applied in previous work (64), ensuring the consistency of the IDL’s H or the CC-site across the ensembles while capturing the structural flexibility and disorder inherent in the flanking regions. To create the monomer, dimer and trimeric states of N_1-246_, we extended the linker to encompass the NTD and the disordered Narm regions. For N_1-365_, we appended the CTD, and for N_1-419_, we included the disordered Carm region as well. The Narm and Carm segments were modeled with FM, while NTD (PDB: 7CDZ) (65) and CTD (PDB: 6ZCO) (66) were constrained to their solved X-ray structure, following approaches used in previous studies (67–69). For each built segment, side-chains were added using SCCOMP (70) and subsequently pre-processed with Rosetta 3.5 fixbb-module to alleviate steric clashes. The final models were energy-minimized with GROMACS 5.1.1 (46).

Each ensemble comprised 10,000 conformers per oligomeric state, representing a diverse sampling of possible structural configurations for monomeric and self-assembled states (dimer, trimer or tetramer). SAXS datasets were analyzed using the theoretical SAXS profiles back-calculated for each conformer in the ensembles. All theoretical curves were computed using CRYSOL (62) with 101 points and a maximum scattering vector of 0.5 Å^-1^ using 25 harmonics. The Ensemble Optimization Method (EOM) was applied to the generated ensemble models to select sub-ensembles that best fit the experimental SAXS profiles. The goodness-of-fit of these selected sub-ensembles was evaluated using the χ^2^ test, calculated as

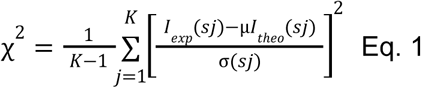

Where *K* is the number of data points in the SAXS profile *I*_*exp*_ (*sj*), σ(*sj*) is their standard deviation, and µ is a scaling factor. The theoretical SAXS curve *I*_*theo*_ was obtained by averaging the scattering of the selected models (*N*_*se*_) as:

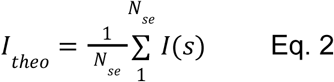

Here, *N*_*se*_ represents 50 explicit models from each EOM run. Representative sub-ensembles for each oligomeric state and construct were deposited in the Protein Ensemble Database (PED) (71), with identifier codes provided in **Supplementary Table 5**.

From multiple independent runs, the structural metrics for the EOM-derived ensembles were calculated using MDAnalysis (50) and ATSAS (62) with python *in-house* scripts as described below.

End-to-end-distance (EED) and Volume for each ensemble member were calculated using the ffmaker module from the ATSAS software suite and the Kernel density estimation (KDE) plots were performed using the SciPy python as implemented in the Seaborn python library.

The centre-of-mass (COM) distance distributions were calculated using MDanalysis by taking the euclidean distance between the indicated structural elements with the KDE plots being calculated as described above. The 3D distributions of the COM distances were calculated as described previously (69), and visualised using Compiled Graphics Objects (CGO) as implemented in Pymol (72).

### Turbidity assays

Turbidity was used to evaluate the effect of ionic strength on N-protein phase separation. To this end, GFP-fused full-length N-protein and its L3P mutant were purified with 500mM NaCl and diluted to 10µM in a buffer containing 100mM Tris·HCl pH=7.2, 5mM *β*-mercaptoethanol, 1mM EDTA at the indicated NaCl concentrations (10, 25, 50, 100, 150, 200, 275, 350, 425 and 500mM) in triplicates on 96-well plates with 50 μL samples. The LLPS phenomena were monitored by measuring the 340nm absorbance in each well every 10 min for 140 min in an absorbance microplate reader (Biotek) and subtracted from blanks without protein. Time-dependent turbidity was fitted to a exponential decay (Eq. 3):

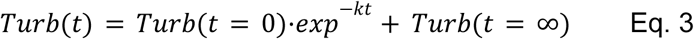

where *Turb*(*t*) is the turbidity along time; *Turb*(*t* = 0) is the turbidity in the initial time (*t* = 0); *k* corresponds to the decay rate constant; *t* is time (in minutes); and *Turb*(*t* = ∞) is the turbidity when *t* = ∞. Estimated parameters are in **Supplementary Table 6**.

### Confocal microscopy *In vitro* Phase Separation

GFP-fused N_1-419_ wild-type and L3P variants were analysed by fluorescent microscopy, monitoring droplet formation. Purified proteins were diluted to 10 μM in a buffer containing 100mM Tris·HCl, 5mM beta-mercaptoethanol, 1 mM EDTA, pH=7.2, at the indicated NaCl concentrations (25, 150, and 250 mM). Sample preparation was done by placing 4 μL onto a glass slide and measuring ca. 15 min after deposition.

We acquired images in triplicate on a Zeiss LSM 880 point scanning confocal microscope controlled with the Zeiss Zen 2.3 (black edition) software, using a gallium arsenide phosphide (GaAsP) PMT detector, a 20x/0.8 Plan-Apochromat differential interference contrast (DIC) Air objective and the 488 nm laser line. Image size for subsequent analysis was 425.10 μm. We used the Fiji software (73), for droplet size calculations and relied on its segmentation feature. We adopted the OTSU threshold and selected droplets based on a predominantly circular shape (over 90% circularity) in an ellipsoid approximation. Total number of fitted ellipses used for distribution plots was 4456, 3691, 3348 and 783 for N_1-419_ wild-type at 25 mM NaCl, N_1-419_ wild-type at 150 mM NaCl, N_1-419_L3P 25 mM at NaCl and N_1-419_L3P 150 mM at NaCl, respectively. The Mann-Whitney U-test assessed the differences between droplet radius distributions, as reported by the major radius of the fitted ellipse (74).

We used fluorescence recovery after photobleaching (FRAP) to investigate the dynamics of GFP-fused N_1-419_ wild-type and L3P variants within the droplets. Purified proteins were diluted to 10μM in the previous buffer at 25 mM NaCl. For those experiments, circular regions slightly smaller than the droplet border were chosen as regions of interest (ROI) and bleached after 5 scans with 100 iterations at full laser power. Recovery was imaged at 1% laser intensity to a total of 500 frames with 770 ms of interval. The fluorescence intensity for each ROI was corrected by subtracting a background region where no fluorescence was detected. FRAP curves were normalised by setting the pre-bleach fluorescence intensity values to 1 and the intensity immediately after bleach to zero. We fitted a mono-exponential function for the case of N_1-419_ wild-type (14) :

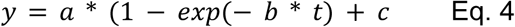

where b is the time constant and (a + c) the maximum value at time (t)= ∞, corresponding to the mobile fraction.

For N_1-419_L3P, a bi-exponential function provided a better fitting (14):

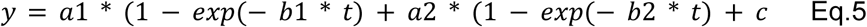

a1 and a2 are the individual maximum signal values of each component, and b1 and b2 are the time constants of the two exponentials.

## RESULTS

### The N-protein interdomain linker has a central amphipathic helix

The SARS-CoV-2 N-protein is a multidomain protein with a hybrid globular/disordered architecture (∼52% disorder) conserved among β-coronaviruses and SARS-CoV variants (**Figure 1A**). The analysis of multiple sequences shows that disordered regions flank (Narm and Carm) and connect (IDL) both folded domains (NTD and CTD). Notably, the central IDL, yet largely disordered, displays a conserved sequence with a high propensity for an α-helical structure among variants around residues 215-240 (**Figure S1A**). This region shows a low variability (Shannon entropy) (**Figure 1A**) and is conserved for variants of interest (VOI) (**Figure 1B**). It exhibits a heptad of hydrophobic and polar residues denoted ***abcdefg*** (**Figure 1B**). This pattern often encodes for amphipathic α-helices, which assemble into coiled-coils (CCs) to bury the hydrophobic surface formed by the residues in positions ***a*** and ***d*** (***75***). To assess its helicity, we measured the ellipticity from a peptide harbouring the central residues 215-240 of SARS-CoV-2 N-protein (N_215-240_). The circular dichroism (CD) profile of N_215-240_ has positive values below 200 nm and two negative bands at 208 and 222 nm, commonly associated with well-structured secondary conformations, mainly α-helical (**Figure 1C**). Its thermal denaturation shows a progressive loss of stable helix structure (**Figure 1C, Figure S1B**). Contrary, the band at 222 nm becomes progressively more negative with peptide concentration, revealing an increase in helicity (**Figure 1D**) and θ_222_/θ_208_ ratios > 1.0 (**Figure 1D, Figure S1B**) that support α-helical coiled-coil structure. Moreover, AlphaFold-2 (AF-2) (35, 38) predicts a well-defined coiled-coil dimer (CC-Di) for N_215-240_ (**Figure S2A**) with high confidence and low interchain uncertainty (**Figure S2B, C**). Conserved leucines mediate the CC-Di interface, further stabilized by electrostatic interactions through ***e*** and ***g*** residues, i.e., the electrostatic pair R226-E231 (**Figure S2A, B**).

**Figure 1.**
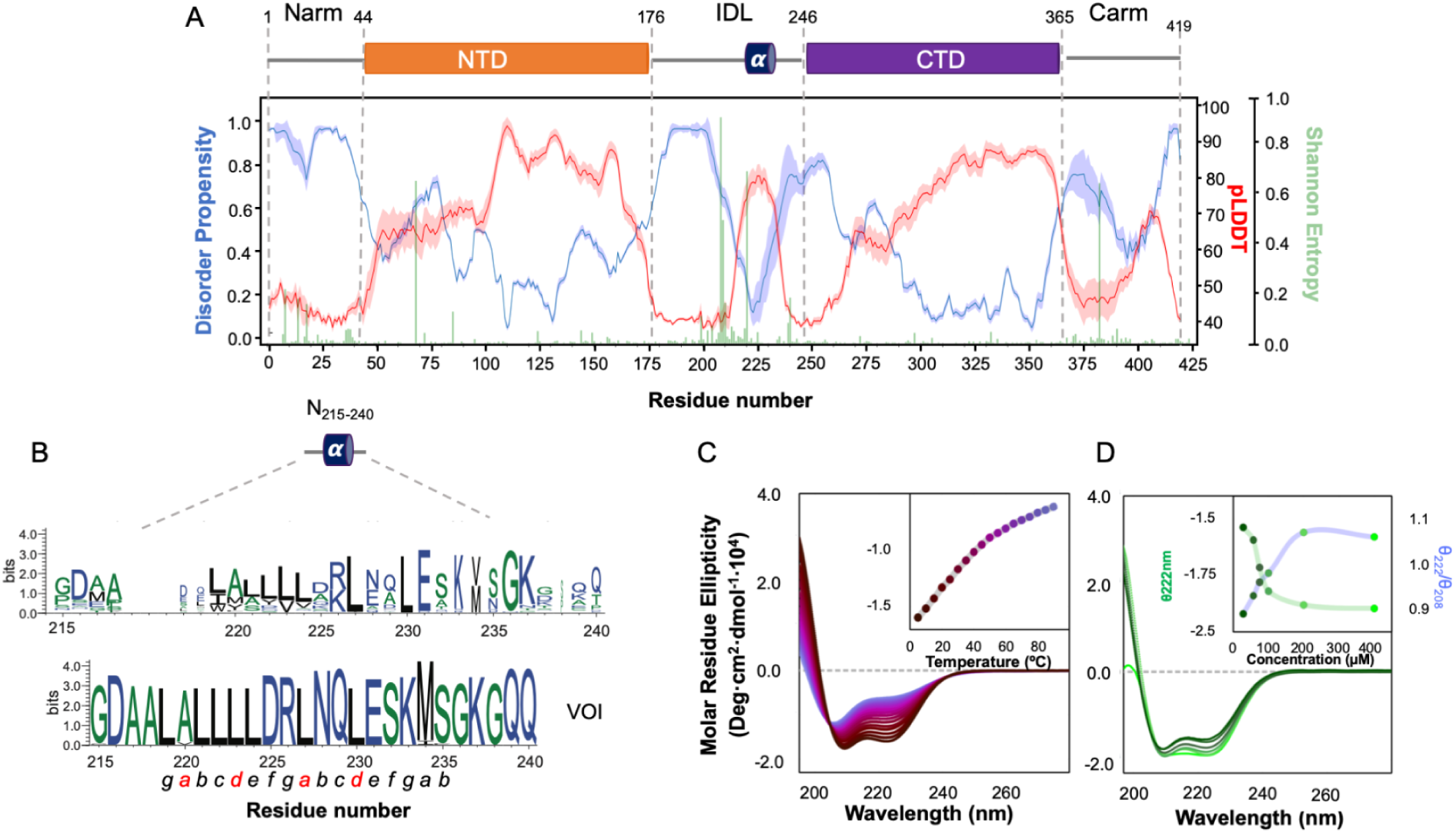
N-protein disordered linker bears a conserved coiled-coil site. (**A**) N-protein conserved globular/disordered architecture. Disorder propensities were calculated with Metapredict (blue) (28) and predicted structure confidence from AF-2 (35) (red) for N-protein variants. Shaded areas correspond to the 95% confidence interval of the corresponding metric. Both predictions, mutually in anti-phase, agree with the organisation of two folded domains flanked by disordered termini and a linker that displays a short central region with low disorder and high AF-2 confidence. The green bars are the per-residue Shannon entropy. On top is the domain schematic of the SARS-CoV-2 N-protein, with domain boundaries marked by dashed lines and NTD and CTD in orange and purple boxes. A blue cylinder respects the region with low disorder and high AF-2 confidence. (**B**) Sequence Logo (30) showing the most conserved residues for the mid-segment N_215-240_ within IDL (top). This central region is conserved among VOI (middle). Heptad pattern in which the *a* and *d* positions in red are occupied by hydrophobic residues (bottom). (**C**) Far-UV circular dichroism spectra of N_215-240_, revealing the presence of α-helical structure. **Inset:** Thermal denaturation followed by the changes in the CD signals at 222 nm, recorded from 5 to 95°C. (**D)** Far-UV CD spectra at 5°C were recorded at different concentrations. **Inset:** Mean Residue Ellipticity at 222 nm (purple line) and ratio θ222/θ208 (light green line) evolution against peptide concentration, reporting the likelihood for the α-helix being in isolation or found within a coiled-coil structure.

N-proteins from β-coronaviruses have a shared architecture and substantial sequence similarity (**Figure 1A**), including the leucine-rich central α-helix within the IDL, thus highlighting functional importance. Leucine-rich regions are found in many proteins and are often involved in protein-protein interactions. Moreover, α-helical coiled-coils are often versatile subunit oligomerization motifs in proteins (76). So, our observations support that the IDL of N-protein has an amphipathic α-helix capable of coiled-coil-driven self-assembly, which we assess in this work with structural detail. To further characterise this assembly, we turned to molecular dynamics (MD) simulations, nuclear magnetic resonance (NMR), small-angle X-ray scattering (SAXS), and mutation analysis.

### N-protein interdomain linker self-assembles by parallel coiled-coil interactions

AF-2 predicts that residues N_215-240_ form a parallel CC-Di packed by *knobs-into-holes* interactions (41), involving residues A220, L223, L227, L230 and M234. These residues create a hydrophobic patch along the peptide helix axis. Although this sequence could theoretically support an antiparallel topology (**Figure S3A**), our two 1.5 μs unbiased all-atom MD simulations indicate that the parallel topology is more likely to occur. In the simulations, the parallel topology undergoes smaller structural displacements compared to the antiparallel model, which displays higher root-mean-square fluctuations (RMSF) (**Figure 2A**). While both models share similar residues at the CC-Di interface, the parallel topology shows more stable inter-chain contacts over time (**Figure 2B**). Native contact analysis reveals that the parallel dimer retains 60-80% of the initial inter-chain contacts, whereas the antiparallel dimer loses 80% of these contacts within 0.8 μs (**Figure 2B**). Thus, MD supports that for N_215-240_, the parallel topology is more plausible *in silico*, with L223, L227, and L230 being critical, exhibiting higher contact frequencies than the antiparallel model (**Figure 2C, D**).

**Figure 2.**
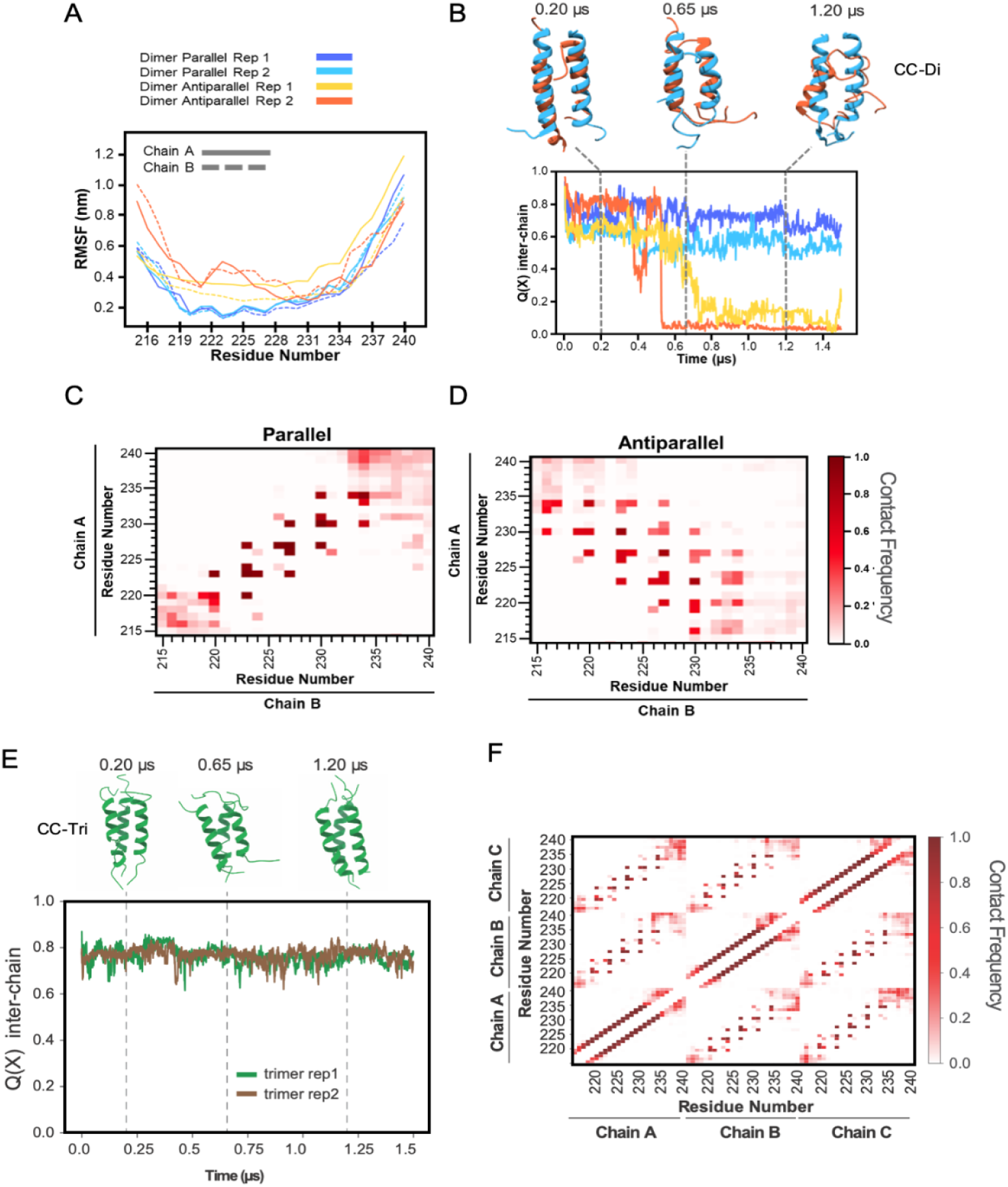
N_215-240_ parallel coiled-coil site. (**A**) Root mean square fluctuation (RMSF) from molecular dynamics of the parallel and antiparallel dimer states. Solid and dashed lines represent chains A and B, respectively. Rep1 and rep2 are trajectory replicas 1 and 2 of 1.5 μs, respectively. (**B**) Inter-chain native contacts for the dimer states referred to in panel C. Colour codes are identical to panel C. Representative snapshots for the trajectories, retrieved from the first replica, at 0.2, 0.65, and 1.2 μs have the same colour codes as in panel A. (**C**) Average contact maps for the parallel dimer and (**D**) antiparallel states retrieved from the trajectories. **(E)** Inter-chain native contacts for the trimeric state. In green and brown are the first and second replicas, respectively. Representative snapshots, in green, for the first replica are presented in cartoon representation for the timestep as in panel B. **(F)** Average contact map for the trimer state.

Recent studies have shown that residues 210-246 of SARS-CoV-2 N-protein, encompassing the N_215-240_ segment, can assemble into trimeric coiled-coils (25). Moreover, N_215-240_ contains a trimerization motif (**Figure S3B**) that specifies a three-stranded, parallel topology in short coiled-coils (27). This specific trimerization motif (RhxxhE) might enhance the propensity of N_215-240_ to form a stable trimeric coiled-coil structure (CC-Tri). Consequently, we extended our MD analysis to explore the CC-Tri configuration. The results demonstrate a stable CC-Tri interface (**Figure 2E**) involving leucines L223, L227, and L230 over time (**Figure 2F**). In agreement with our simulations, we observed experimentally that those residues are indeed at the CC interface.

We assigned the backbone resonances of a construct encoding the IDL sequence (N_176-246_) (up to 95.8%), including the missing resonances within the 215 to 240 region not observed in previous work (**Figure 3A**, **Figure S4**) (77, 78). This achievement provided access to high-resolution information about the CC interface. Figure 3A shows the ^1^H–^15^N heteronuclear single quantum coherence (HSQC) NMR spectra of IDL across varying concentrations. These spectra exhibit low dispersion of the ^1^H chemical shifts, as expected for an IDR (**Figure S4A**). However, the structural propensity derived from the NMR chemical shifts (79) supports the tendency of N_215-240_ to adopt a helical conformation (hereafter helix H), with the rest of the IDL disordered (**Figure 3B**). Consistent with self-assembly within IDL, we observed detectable peak shifts (**Figure 3A, C**) and differential line broadening leading to a concentration-dependent intensity drop (**Figure 3A, D**) for the resonance around residues 215 to 237. The observed intensity drop displays an *i+3* / *i+4* pattern oscillation, compatible with coiled-coil interactions matching the AF-2 model interface residues leucine 223, 227 and 230, and sustained by MD. Moreover, the helical region shows relatively elevated *R_2_* values (**Figure 3E**) that increase with concentration, which, together with the observed broad resonances, are consistent with structural heterogeneity and conformational exchange between monomers (80). Overall, NMR thus captures, with site-specific resolution, a CC site in the IDL of the SARS-CoV-2 N-protein driving self-assembly.

**Figure 3.**
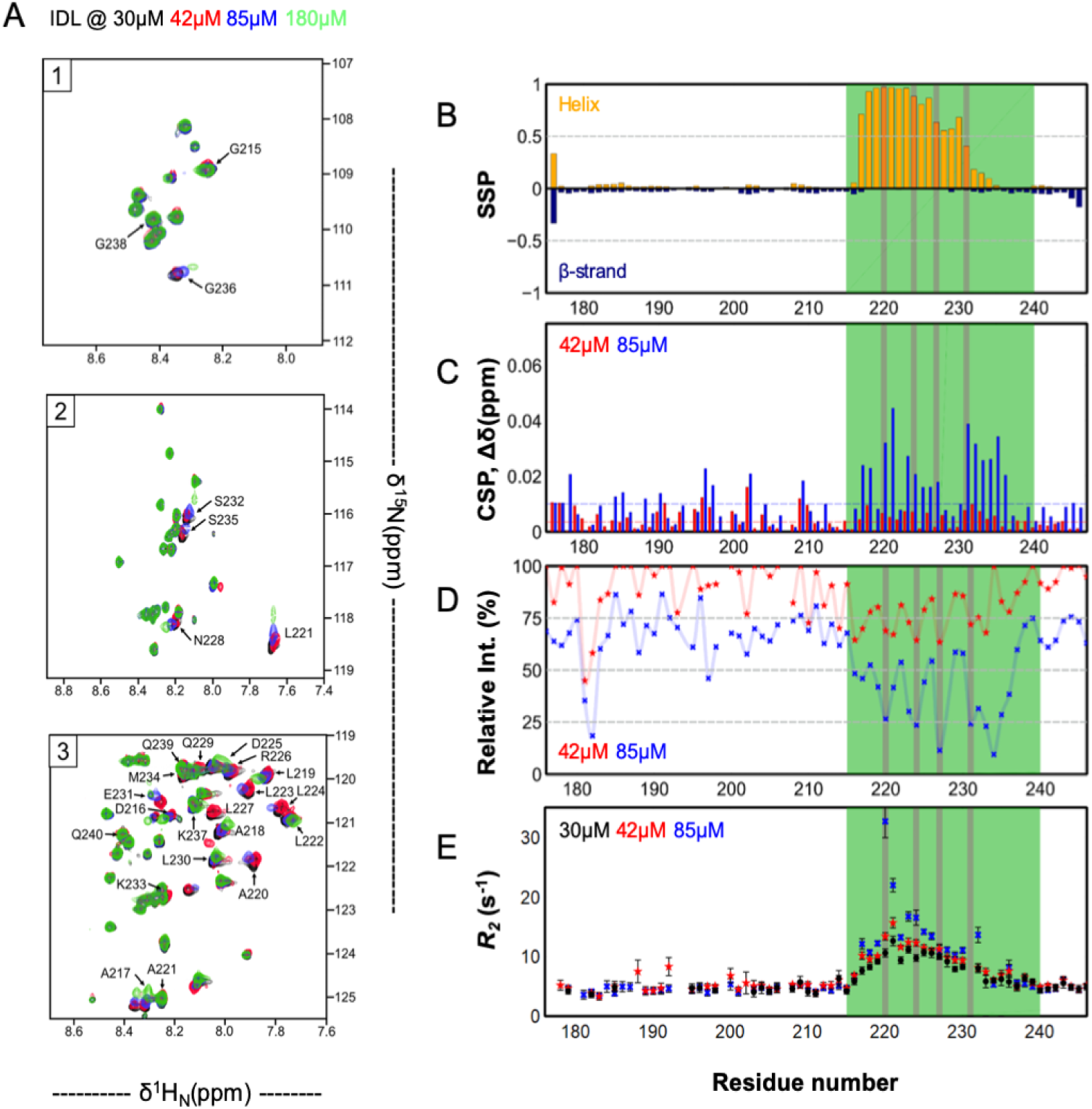
Capturing N_215-240_ coiled-coil self-assembly with NMR. (**A**) Expansions of [^15^N-^1^HN]-HSQC spectra of IDL (N_176-246_) in **Fig. S4** at different concentrations, showing the broad resonances observed due to concentration-dependent self-assembly. Resonances of N_215-240_ are labelled (1-letter code and number) in the plot. (**B**) Secondary structure propensity for β-strands (blue) and helices (orange) calculated from protein backbone chemical shifts at 42µM. Residues within the region 215-240 (green shading) display high helical propensity. (**C**) [^15^N-^1^HN]-chemical shift perturbations (CSP) from the reference spectrum at 30uM plotted as a function of the sequence of IDL. (**D**) Site-specific NMR signal attenuation is observed for the IDL resonances, affecting mainly the α-helix, with the resonances of the *knobs-into-holes* residues becoming broad and exhibiting the *i+3* (220, 223, 227, 230) and the *i+4* (223, 227, 230, 234) patterns of helical conformation. The vertical bars (purple shading) denote the position of the *knobs-into-holes* (i.e., *a, d*) residues. (**E**) ^15^N–*R_2_* analysis of IDL at different concentrations. Relaxation rate errors were derived from the covariance method of the residuals’ fitting.

### Dynamic Monomer-Dimer-Trimer Ensemble of the N-Protein Interdomain Linker

To elaborate further on our previous observations, we engineered a mutant (L3P) by replacing leucines 223, 227, and 230 with prolines to disrupt the helical structure of H and coiled-coil interactions (**Figure 4A)**. We then monitored the self-assembly process using SAXS.

**Figure 4.**
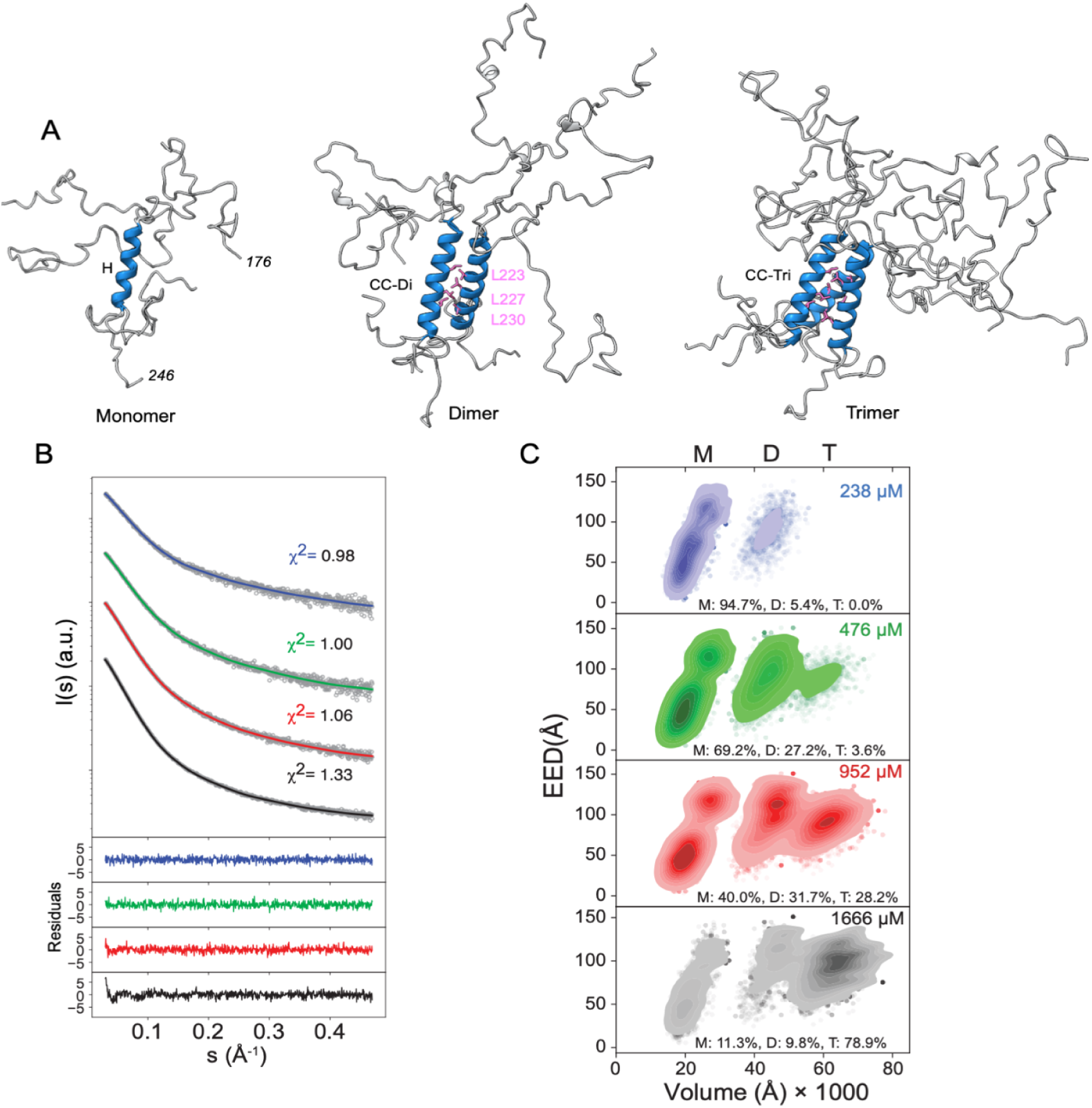
Interdomain linker monomer-to-trimer ensembles. (**A**) Representative sub-ensembles of monomer (M), dimer (D) and trimer (T) states of N_176-246_ (IDL) are depicted in cartoon representation, highlighting the 3 top-selected conformers per state. The helical region H is emphasized in the IDL monomer ensemble. Leucines 223, 227, 230 are shown in pink stick representation in both CC-Di (dimer) and CC-Tri (trimer) models. **(B)** Experimental SAXS profiles for N_176-246_ at different concentrations are shown in grey and respective EOM fittings in solid lines. As a figure of merit, we use the χ^2^ test to assess the goodness-of-fit between the experimental data and each sub-ensemble model. Point-by-point residuals of the fittings are displayed at the bottom. (**C**) Kernel density contour plots for End-to-End distance (EED) and Volume, calculated from the EOM-selected sub-ensembles, are provided next to the SAXS curves panel and colored using the same color code. “M”, “D” and “T” denote monomer, dimer and trimer species, respectively.

To explore the conformational landscape of monomer to coiled-coil self-assembled states of the N-protein IDL in solution and its dependence on CC interactions, we acquired SAXS data on N_176-246_ WT and N_176-246_L3P (CC-null) (**Supplementary Tables 3 and 4)** across varying concentrations. The SAXS data for N_176-246_ WT reveals a concentration-dependent evolution consistent with self-assembly dynamics (**Figure S5A**). This contrasts sharply with the SAXS profile for the N_176-246_L3P. At lower concentrations, the SAXS-derived Kratky plots for both constructs exhibit a gradual increase in (*sRg)2·I(s)/I*(*0*) at wide angles, characteristic of highly disordered proteins (**Figure S5B**) (81). As concentration rises, N_176-246_ WT manifests a broad peak, indicating the emergence of structural organization. Conversely, the N_176-246_L3P mutant maintains a disordered profile that remains largely unaffected by changes in concentration.

Quantitative analysis of SAXS-derived *Rg* and *D_max_* parameters provides additional insights. The L3P mutant displays relatively stable *Rg* and *Dmax* values, suggesting minimal alterations in conformation with changing concentrations (**Figure S5C**). In contrast, the wild-type IDL displays an increase in these parameters at higher concentrations, plateauing around 45 Å and 190 Å in *Rg* and *Dmax*, respectively.

To comprehensively understand IDL’s conformational properties and oligomeric states, we generated extensive ensembles representing monomeric, dimeric, and trimeric states (**Figure 4A**) using the Flexible-Meccano (FM) algorithm (63) combined with AF-2. Employing the Ensemble Optimization Method (EOM) (82) on this pool of structures and self-assembled states enabled the selection of sub-ensembles that faithfully reproduced the scattering profiles (**Figure 4B**). For the IDL wild-type, we obtained 94.7% monomeric conformers (X^2^=0.98) from its lowest concentration data (ca. 238µM), contrasting with the 78.9% of parallel trimer conformers for the highest concentration measured (ca. 1666µM) (**Figure 4C**). As the concentration increased, the population of self-assembled states also increased, indicating concentration-dependent assembly dynamics. Emphasizing the pivotal role of the helical region (H) in self-assembly, all SAXS curves for the CC-null mutant (N_176-246_L3P) nearly exclusively yielded monomeric conformers (**Figure S6A, B**). Despite transitioning to self-assembled states, the SAXS-based ensemble revealed that the IDL dimer and trimer remain highly flexible and disordered.

To further validate the self-assembly state, we extended the linker to encompass the NTD and the disordered Narm regions (**Figure 1A**), creating the N_1-246_ construct, which we then probed using SAXS. The resulting data was analyzed using explicit ensemble models of monomers, dimers and trimers. These models were built using the X-ray structure of NTD (PDB:7CDZ) (65) and modeling the disordered regions of the IDL and Narm with FM (**Figure 5A**).

**Figure 5.**
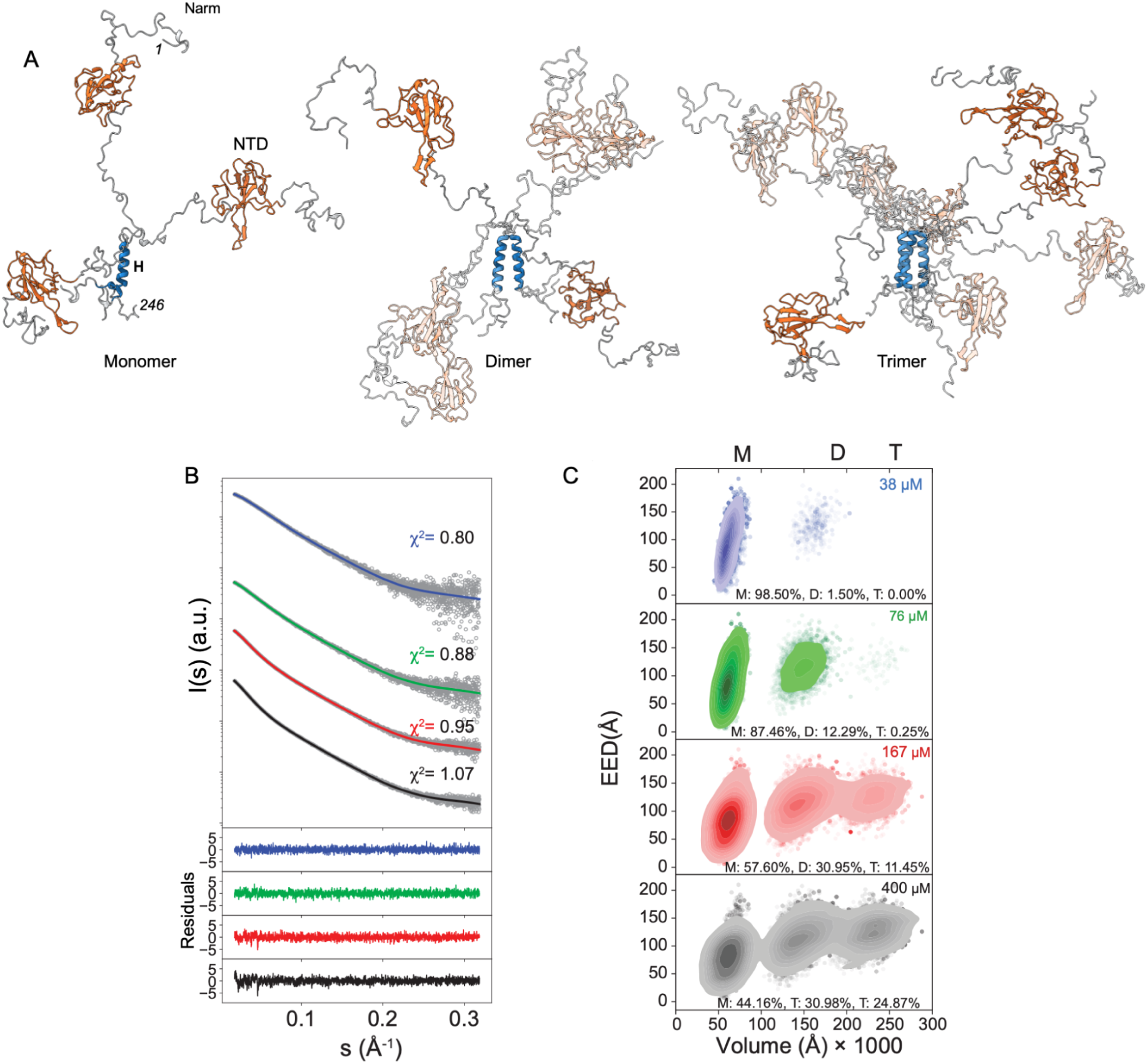
N_1-246_ monomer-to-trimer ensembles. **(A**) Representative sub-ensembles of explicit monomer and coiled-coil dimer and trimer states of N_1-246_ are depicted in cartoon representation, including the 3 top selected conformers. The helical region H, the NTD and Narm are indicated in the N_1-246_ monomer ensemble. **(B)** Experimental SAXS profiles for N_1-246_ at different concentrations in grey and respective EOM fittings in solid lines. As a figure of merit, we use the χ^2^ test to assess the goodness-of-fit of each sub-ensemble. Point-by-point residuals of the fittings are at the bottom. (**C**) Kernel density contour plots for End-to-End distance (EED) and Volume, calculated from the EOM-selected sub-ensembles, are provided next to each SAXS curves panel and colored using the same color code. “M”, “D”, and “T” denote monomer, dimer and trimer species, respectively.

This analysis revealed that N_1-246_ coexists in a monomer-dimer-trimer equilibrium in solution, indicating a dynamic behaviour similar to that observed for the N-protein IDL. Notably, the presence of Narm and NTD does not significantly alter the oligomerization state of N_1-246_, suggesting that the factors driving the oligomeric equilibrium are likely intrinsic to H sequence or structure rather than being strongly influenced by the presence of additional domains. At higher concentrations (ca. 400 μM), self-assembled states become more prominent, with the trimeric form representing 24.87% of the population (55.85% self-assembled states). At lower concentrations, SAXS predominantly probes the monomer (98.5% at ca. 38 μM) (**Figure 5B, C**).

In contrast, the N_1-246_ CC-null variant (N_1-246_L3P) shows a much lower proportion of dimers (8.97% at 295 μM), with no significant trimer formation (**Figure S6C, D**). Additionally, the self-assembly is reflected in an overall shift of the *P(r)* distribution towards larger distances for N_1-246_ WT, indicating larger conformations (**Figure S7A**). This shift is less pronounced for the N_1-246_ L3P mutant, both in *Rg* and *Dmax* (**Figure S7B**). These findings support the notion that the coiled-coil structure significantly contributes to the overall stability of self-assembled states. This ensemble analysis also captures the conformational sampling of NTD while tethered to the pre-formed helix within the IDL. This structural plasticity is evident in the ensemble’s representation of inter-domain distances (**Figure S8A**) covering a broad distribution range (**Figure S8B-D**). The distance distributions between the CC site and NTD (D_CC-NTD_), inter-chain NTDs (D_NTD_), and *Rg* distributions for the N_1-246_ EOM-selected sub-ensembles show more extended distributions compared to the random ensemble. These deviations are less pronounced for the monomer state (**Figure S8B**) and more for the self-assembled states, with NTDs being 87.33 ± 27.62 Å and 93.6 ± 28.82 Å apart for the dimer and trimer states, respectively (**Figure S8C, D**). The extended conformations of self-assembled states may be functionally relevant, facilitating interactions with other biomolecules. Overall, our comprehensive SAXS analysis further emphasizes the role of the CC site within IDL in directing structural assembly, and not NTD-CC interactions, and the dynamic nature of the self-assembled states, suggesting conformational adaptability in response to environmental cues or interactions (83).

### The linker coiled-coil directs N-protein’s dynamic head-to-head high-order structures

SAXS effectively studies the spatial distribution of globular domains within flexible proteins, capturing the polydispersity of transient and multivalent macromolecular assemblies and oligomers (69, 84–87). Leveraging SAXS’s ability to probe these dynamic systems, we investigated the conformational and oligomeric states of constructs encompassing the CTD (N_1-365_) and the Carm (N_1-419_) (**Figure 1A)** (**Supplementary Table 2)**. These proteins were expressed, purified, and analyzed at varying concentrations using explicit ensemble models of the oligomeric species.

SAXS data for N_1-365_ and N_1-419_ revealed a concentration-dependent self-assembly, akin to that observed for N_176-246_ and N_1-246_. This is evidenced by variations in their *P(r)* functions and Kratky plots (**Figure S9A, B)**. The *P(r)* curves (**Figure S9A)** are highly asymmetric and bimodal, with two distinct peaks reflecting the intra- and interdomain pairwise distances. With increasing concentrations, the interdomain distance peak becomes more pronounced, followed by increases in *Rg* and *Dmax* (**Figure S9C**), indicating progressive assembly. The Kratky plots (**Figure S9B)** deviate from the typical bell-shaped of globular, consistent with the flexible, multidomain nature of these constructs (88).

The N-protein forms highly stable dimers via its CTD (65, 66), driven by an antiparallel swap dimerization of the CTD (**Figure S10**). This interaction, referred to as **site 1** or the “**tail**” (**Figure 6A**), arranges the linker segments connected to helix H outward, with anchor points approximately 40 Å apart on each monomer (**Figure S10**). Additionally, based on the self-assembly equilibrium observed for N_176-246_ and N_1-246_, the CC site within the IDL emerges as a secondary self-assembling region (**site 2**, or the “**head**”), capable of facilitating either intra-dimer interactions (closed state) or inter-dimer interactions (open state), enabling higher-order structures (**Figure 6A**).

**Figure 6.**
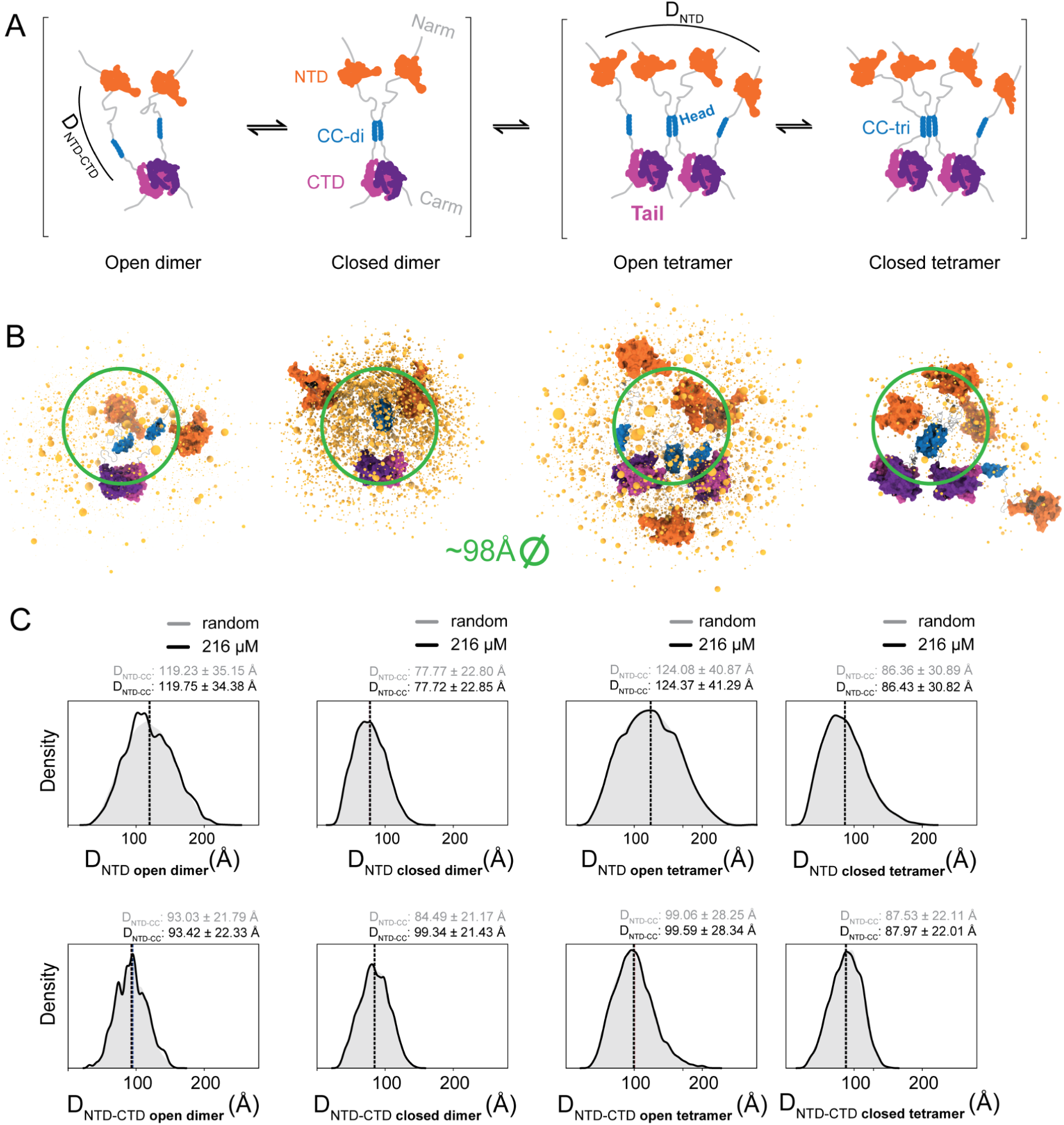
Conformational landscape and structural metrics for the dimer-tetramer equilibrium of N_1-419_. (**A**) Schematic representation of N_1-419_ coiled-coil (CC)-driven oligomerization. The assembly includes open and closed dimeric and their equilibrium with tetrameric states. The CC motif (blue) facilitates parallel, head-to-head and tail-to-tail oligomerization of N-protein dimers. The N-terminal domains (NTDs) are in orange, the C-terminal domains (CTDs, tail) are in purple and light purple, and the inter-domain linker (IDL) along with the N-terminal (Narm) and C-terminal (Carm) tails are in light grey. Distances between NTDs (D_NTD_) and between NTDs and CTDs (D_NTD-CTD_) are indicated. CC-di and CC-tri are the dimer and trimer coiled coils, respectively. (**B**) Conformational landscape at 216 µM, illustrating the open and closed dimeric and tetrameric states. Structures are displayed using the same colour scheme as in panel A, with NTD, CC, and CTD domains represented as surfaces, and the IDL in cartoon representation. Centers-of-mass for NTDs, derived from the EOM-selected SAXS ensemble at 216 µM, are depicted as yellow spheres. The size of each sphere reflects the frequency of the corresponding conformation within the ensemble (larger spheres indicate higher probabilities). A green circle provides a reference for scale. (**C**) KDE plots for D_NTD_ and D_NTD-CTD_ distance distributions based on the EOM-selected SAXS ensembles of open and closed dimeric and tetrameric states at 216 µM. The black KDE curves represent the selected ensemble, while the filled grey areas denote the KDE distributions of corresponding random ensembles for comparison.

To investigate the role of the CC interactions at site 2, we modelled both “open” and “closed” dimers, where IDLs are either spatially separated or interacting through site 2, respectively. Full-atom ensemble representations were further created for plausible tetramer states driven by the formation of CC-di or CC-tri structures at site 2 (**Figure 6A**). SAXS data analysis revealed the relative populations of these states, showing that at low concentrations, closed dimers predominate, while at higher concentrations, open dimers and tetramers become increasingly prevalent (**Figure S11A, S12A)**.

Explicit ensemble models (**Figure 6B**) derived from SAXS profiles (**Figure S11B, S12B**) validated these assemblies, supported by robust χ^2^ metrics. Notably, excluding open states significantly worsened the goodness-of-fit (χ^2^) to the SAXS data (**Figure S11C**), corroborating the open-to-closed equilibria in dimer and tetramer assemblies. The closed dimer is characterized by more compact conformations stabilized by CC interactions at site 2, while the open dimer represents a precursor to tetramers, with more spatially separated IDLs (**Figure 6B**).

Tetrameric states primarily adopt two configurations: symmetric (open) tetramers, with flexible IDLs interacting via CC-di, and asymmetric tetramers (“closed”), involving CC-tri and one unengaged monomer at site 2 (**Figure 6A**, **Figure 6B**). The SAXS-derived conformational landscape indicates that open tetramers dominate at higher concentrations, underscoring the role of the IDL’s CC motif in mediating higher-order assembly.

Our findings also show that the N_1-419_ assemblies occupy a broad conformational space (**Figure 6B, 6C, Figure S11D**), with interdomain distances resembling those of random coils and remaining largely concentration-independent. This indicates that oligomerization is primarily driven by CC interactions at site 2. Similar behaviours observed for N_1-365_ (**Figure S12C**) reinforce these conclusions across constructs. Intradimer interactions mediated by the IDL’s CC motif play a significant role in shaping the spatial distribution of the NTD, thereby influencing its structural and functional dynamics. Both dimeric and tetramic states transition between compact (closed) and extended (open) configurations, modulated by the interaction network within the IDL. Open dimers and tetramers exhibit an average D_NTD_ interdomain distances of 119.75±34.38 Å and 124.37±41.29 Å, respectively, while their closed counterparts show distances of 77.72±22.85 Å and 86.43±30.82 Å (**Figure 6B**). These structural transitions between open and closed states, along with the dynamic equilibrium of oligomeric forms, are likely central for N-protein’s diverse roles, including RNA genome organization, replication, and efficient viral assembly. Interestingly, the relative spatial distribution between the NTD and CTD, reflected by the D_NTD-CTD_ metric, remains largely unaffected by these CC-driven transitions (**Figure 6B**). This spatial separation likely allows the N protein to simultaneously engage in RNA binding and the formation of higher-order assemblies, processes critical for efficient viral genome packaging.

### The CC interdomain helix modulates N-protein phase separation

N-protein of SARS-CoV-2 undergoes LLPS to form biomolecular condensates, a process crucial for various aspects of the virus’s lifecycle. Given the significance of the IDL central helix in N-protein homo-oligomerization, we aimed to investigate its role in modulating LLPS. We conducted turbidity assays and confocal microscopy experiments to assess the impact of mutations abolishing the CC site within the IDL on the stability and kinetics of LLPS under different salt conditions.

Electrostatic interactions play a pivotal role in the LLPS of N-protein, with purified N-protein in high salt concentrations unable to form droplets with RNA (89). Therefore, we first monitored the turbidity of full-length N-protein and N_1-419_L3P (a mutant lacking the CC site) at various salt concentrations over time to probe their LLPS behaviour. LLPS was induced by diluting the protein from a solution of 500 mM NaCl into a buffer with lower salt concentrations. Our results indicate that both N-protein variants can undergo LLPS, without added RNA, via homotypic interactions (**Figure 7**). Moreover, as observed for N:RNA droplets (89), ionic strength also challenges their LLPS process.

**Figure 7.**
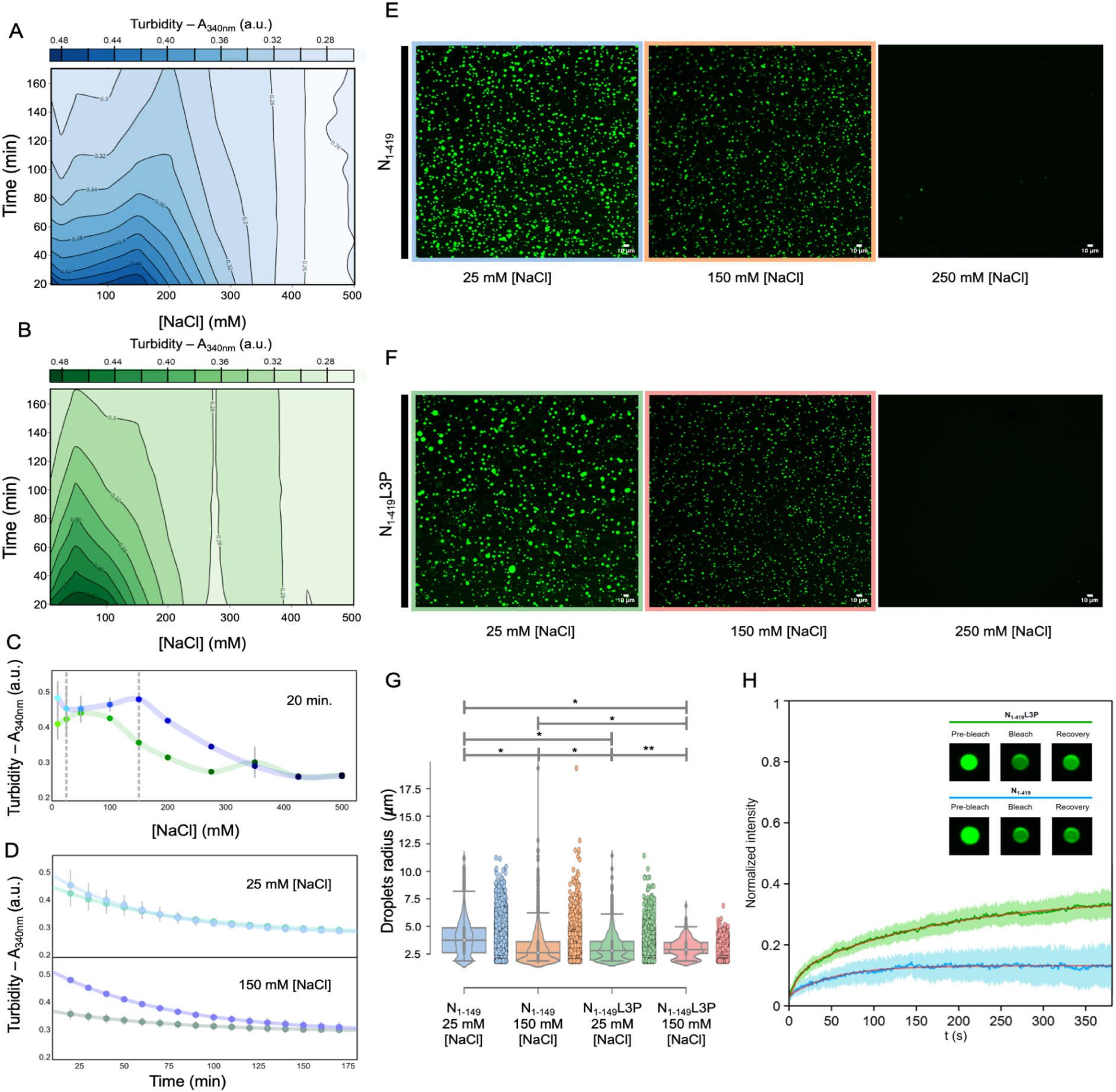
Impact of CC Interdomain Helix on N-Protein LLPS. (**A, B**) Contour plots illustrating the turbidity at 340 nm of N_1-419_ (depicted in blue gradient) and N_1-149_L3P (depicted in green gradient) over time at various salt concentrations, highlighting the influence of the CC interdomain helix on LLPS. (**C**) Turbidity signals after 20 minutes of incubation at different NaCl concentrations for N_1-419_ (blue dots) and N_1-419_L3P (green dots), with dashed lines indicating 25 mM and 150 mM conditions. (**D**) Turbidity profiles over time for N_1-419_ (blue dots) and N_1-419_L3P (green dots) at 25 mM (top) and 150 mM NaCl (bottom) showcasing the differential behaviour of the two variants under varying ionic strengths. Solid lines are fitting assuming an exponential decay. All turbidity data are mean ± standard deviation (SD) from experiments conducted in triplicate. (**E, F**) Fluorescence images of LLPS droplets formed by GFP-labelled N_1-419_ (top) and N_1-149_L3P mutant (bottom) at room temperature with the indicated concentrations of NaCl. Scale bars, 10 μm. (**G**) Droplet radius distributions presented in box and violin plots alongside respective strio plots for N_1-419_ and N_1-149_L3P droplets imaged at the indicated NaCl concentration. Statistical analysis was performed using the Mann-Whitney U-test. \**p* ≤ 0.000001; **non-significant (ns). (**H**) FRAP analysis of condensates from N_1-149_ (blue) and N_1-149_L3P (green). Data are normalized to mean ± standard deviation (shaded areas) from fluorescence recovery curves (*n*=6 droplets) over time. The red solid lines correspond to the fitting of a mono-exponential and a bi-exponential function for N_1-149_ and N_1-149_L3P respectively. The inset shows representative micrographs of a droplet for N_1-149_ (top) and N_1-149_L3P (bottom) before bleaching, right after bleaching, and at the end of recovery.

Turbidity measurements revealed that condensation was higher at lower salt concentrations and decayed slowly over time, reflecting the formation and maturation of droplets by LLPS (89) (**Figure 7A,B, Figure S13**). Both variants reached maximum turbidity values at low salt concentrations, suggesting efficient LLPS initiation. However, we observed that the LLPS of the wild-type N-protein was more resistant to changes in ionic strength compared to the mutant lacking the CC site. Specifically, at higher salt concentrations (above 150 mM NaCl), the mutant exhibited a lower turbidity signal, indicating increased difficulty in condensation compared to the wild-type N-protein (**Figure 7C**). For both variants, turbidity decreases over time, characteristic of liquid droplets fusing and settling. Interestingly, the mutations did not affect the decay rate, suggesting that the CC site may not directly influence the rate at which N-protein droplets fuse and settle, at least under the conditions tested (**Figure 7D**, **Supplementary Table 6**). These findings suggest that the CC interdomain helix plays a crucial role in modulating N-protein phase separation, influencing its stability under varying salt conditions.

Following the turbidity assays, we used confocal microscopy to visualize the droplets formed by purified green fluorescent protein (GFP)-fused N_1-419_ (**Figure 7E**) and N_1-419_L3P (**Figure 7F**). When observed under the microscope at low ionic strength, GFP-N_1-419_ droplets exhibited flow and fusion throughout the sample (89), displaying a typical spherical shape (**Figure 7E**) with a radius of 3.88±1.51 μm (**Figure 7G**), consistent with previous observations (21). Our examination further revealed a reduction in droplet size and formation at higher salt concentrations (**Figure 7E, F**), aligning with the turbidity assays’ findings. At 250 mM NaCl, the droplet formation is drastically reduced.

Similarly, the purified GFP-N_1-419_L3P mutant also formed microsized droplets that fuse and flow in solutions with 25 mM NaCl (**Figure 7F**), exhibiting a slightly reduced size distribution (3.06±1.15 μm) (**Figure 7G**), compared to the wild type, as evidenced by its smaller *Turb*(*t* = 0) (**Supplementary Table 6**). Furthermore, the LLPS droplets of the mutant also decreased in size and number with increasing salt concentration, ultimately dissolving at 250 mM NaCl (**Figure 7F**). Notably, the circular-shaped droplets formed by the triple mutant exhibited minimal changes in size distribution when the salt concentration was increased to 150 mM NaCl (2.97±0.82 μm). In contrast, the wild type decreased in size under the same salt concentrations, with an average size comparable to that of the L3P variant (2.98±1.4 μm). Additionally, the size distributions of N_1-419_L3P became sharper compared to broader N_1-419_ distributions. These differential behaviour suggest a complex interplay between the CC interdomain helix and other factors influencing droplet size regulation, including electrostatic interactions and other structural features of the N-protein.

FRAP experiments further highlight the significance of the CC site on the LLPS behaviour of N-protein (**Figure 7H**). Consistent with previous studies (14, 17), the wild-type N-protein exhibits slow exchanging dynamics with the soluble pool, achieving a fluorescence recovery of only 13.2±0.2%. This slow recovery indicates stronger self-assembly interactions and stable droplet formation. In contrast, the triple mutant (L3P) shows a more rapid and higher fluorescence recovery (35.7±0.4%), indicative of faster diffusion and a greater mobile fraction. This suggests weakened self-assembly interactions and a less stable, more dynamic phase-separated state.

## Discussion

The N-protein of SARS-CoV-2 is essential to the viral life cycle, facilitating replication, assembly, and interactions with host machinery. Understanding its structural dynamics and functional roles is critical for the development of targeted therapeutic strategies against COVID-19. This study focuses on elucidating the structural and self-assembly properties of the N-protein, particularly its intrinsically disordered linker (IDL), which harbors a conserved coiled-coil (RhxxhE) motif. Our findings reveal critical insights into the oligomerization dynamics and functional versatility of this region, shedding light on its broader implications in viral pathogenesis.

Our analysis identifies a conserved amphipathic α-helix (helix H) within the IDL as a key structural element driving self-assembly. This helix, previously implicated in oligomerization (25), mediates stable intermolecular interactions via leucine residues, forming a parallel CC motif that supports dimer and trimer formation. Using a multidisciplinary approach involving MD simulations, NMR, and SAXS, we confirmed the critical role of this CC motif in driving these oligomeric states and facilitating higher-order assembly. Mutations disrupting CC interactions abolish both dimer and trimer formation, underscoring the motif’s essential role in stabilizing N-protein assemblies.

Our SAXS-based explicit ensemble models reveal that the IDL facilitates a dynamic equilibrium between monomeric, dimeric, and trimeric states stabilized by the CC motif. Constructs lacking the CTD (N_1-246_) exhibit a similar equilibrium, indicating that the IDL and its CC motif are primary determinants of self-assembly, independent of other domains. Moreover, the presence of Narm and NTD does not significantly alter the oligomerization state of N_1-246_, suggesting that this equilibrium is governed by the sequence or structure of helix H rather than being strongly influenced by additional domains. While previous studies have proposed additional transient helices in other regions of the N-protein (13, 23, 90), our findings emphasize helix H as the dominant driver of oligomerization.

In the context of the full-length protein, the IDL facilitates a dynamic equilibrium between dimeric and at least tetrameric states. The CC motif stabilizes these oligomeric forms, mediating transitions between closed (CC motif interacting) and open (CC motifs non-interacting) conformations (**Figure 6A**). At higher concentrations, tetrameric assemblies predominate, indicating that CC interactions serve as a scaffold for higher-order structures. These findings align with previous reports of coiled-coil-driven oligomerization (25). Although the closed (asymmetric) tetramer involving a CC-tri was less populated under our experimental conditions, we do not exclude the potential role of CC contacts forming higher-order assemblies, namely when interacting with RNA, as previously described (91).

Our study identifies a dual role for IDL’s CC motif: stabilizing dimers and mediating inter-dimer interactions to form tetramers, thereby acting as a key structural determinant in N-protein assembly. This head-to-head and tail-to-tail assembly mechanism resembles the organization principles of bacterial H-NS proteins, which facilitate DNA condensation and transcription regulation (92). Such parallels suggest that the N-protein adopts similar structural principles to support its multifaceted functions, including RNA binding, genome packaging, and interactions with host factors.

By facilitating head-to-head interactions through the CC site, the linker emerges as a pivotal structural element that orchestrates the spatial arrangement of NTDs and CTDs, likely crucial for assembling higher-order ribonucleoprotein complexes. The IDL’s conformational dynamics directing theinterdomain spacing appear to influence the structural accessibility of the NTD for RNA binding. This interplay may have functional significance, as the NTD’s ability to bind and recognize RNA structural elements (93, 94), could be modulated by the oligomeric state of the N-protein. The ability of the N-protein to transition between open and closed states likely enables it to adapt its conformation to bind RNA with varying lengths and structures. The flexible interdomain linker may facilitate cooperative binding of RNA by multiple N-protein subunits, enhancing packaging efficiency. Open conformations could allow for better accessibility to RNA binding regions, while closed conformations might stabilize the RNA-protein complex, ensuring proper condensation into ribonucleoprotein particles.

The conserved nature of the CC motif and its role in stabilizing higher-order assemblies suggest that it may also facilitate interactions with other viral or host proteins, such as Nsp3a (23). This interaction with Nsp3a underscores the versatile nature of α-helix H as a motif capable of engaging in both homotypic and heterotypic interactions.

Our findings extend to the role of the IDL in LLPS, a property critical for N-protein’s function in genome condensation and replication compartment formation (17, 24). Mutations disrupting the CC motif alter LLPS dynamics, leading to changes in droplet size and stability. These results reinforce the IDL’s role as a hotspot for modulating LLPS and suggest that environmental cues, such as salt concentration and post-translational modifications (17, 24), could fine-tune phase separation properties. Such regulation may differentiate gel-like condensates promoting nucleocapsid assembly from liquid-like condensates involved in genome processing (24). Additionally, the dynamic self-assembly of the IDL underscores the N-protein’s adaptability, with its architecture and oligomeric states likely modulated by environmental cues (95) or cellular interactions to meet diverse physiological or replication demands. Further investigation into how the CC motif influences intracellular LLPS with RNA could deepen our understanding of viral replication and pathogenesis. For instance, phosphorylation of the SR-rich motif adjacent to helix H may dynamically regulate condensate properties, linking structural modifications to functional outcomes.

Targeting the IDL’s CC motif offers a promising avenue for antiviral intervention. Disrupting its interactions could destabilize N-protein assemblies, impairing genome packaging and replication. Moreover, modulating LLPS properties via small molecules or post-translational modification mimetics (96) may offer additional therapeutic strategies. Future studies should elucidate how the N-protein’s structural dynamics and interactions within condensates influence its role in the viral life cycle.

Through a comprehensive ensemble analysis of N-protein structure and oligomerization, this study highlights the IDL and its CC motif as central to the N-protein’s structural and functional versatility. By unravelling the principles governing N-protein self-assembly and LLPS, we lay a foundation for understanding its role in SARS-CoV-2 biology and pathogenesis. These insights deepen our understanding of coronavirus replication and identify new targets for therapeutic intervention, paving the way for innovative antiviral strategies.

## DATA AVAILABILITY

The NMR chemical shifts for the IDL (N_176-246_) are available in the Biological Magnetic Resonance Data Bank (BMRB) under accession code **52026**. SAXS data are deposited in the Small Angle Scattering Biological Data Bank (SASBDB) (97), with accession codes provided in **Supplementary Table 4**. Structural ensembles for N_176-246_, N_1-246_, N_1-365_, and N_1-419_ have been deposited in the open-access Protein Ensemble Database (PED) (71), with identifier codes listed in **Supplementary Table 6**.

## FUNDING

GH thanks the PT-NMR FCT PhD Program (PD/BD/147227/2019) for financial support. A MOSTMICRO FCT PhD scholarship supports MLM (UI/BD/154576/2022). TNC is the recipient of the CEECIND/01443/2017 grant. National funds funded this work through FCT: Project MOSTMICRO-ITQB (UIDB/04612/2020, UIDP/04612/2020), FEDER Funds through COMPETE 2020 (0145-FEDER-007660), LS4FUTURE (LA/P/0087/2020), and SR&TD project (PTDC/BIA-BFS/0391/2021). Financial support was provided by European EC Horizon2020 TIMB3 (810856). We acknowledge using the ESRF-BM29 (MX-2085-BAG) and DLS-B21 Bio-SAXS beamlines (MX20161-1, SM21035-177, MX25270-8, MX25270-7). NMR data were acquired at CERMAX with equipment funded by FCT, project AAC 01/SAICT/2016. We did microscopy fluorescence experiments at BIC (ITQB NOVA) supported by PPBI (PPBI-POCI-01-0145-FEDER-022122).

## ACKNOWLEDGMENTS

We thank Pedro Matos (ITQB NOVA) for his insightful help and suggestions. We acknowledge Cristina G. Timóteo, Rita Pacheco, Teresa Batista da Silva and João Carita from the Protein Purification and Microbial Cell Production Research Facilities of ITQB-NOVA for their support. We thank Carolina A. Feliciano from Bacterial Imaging Cluster (BIC) for technical support. We thank Ana Catarina Paiva and Tiago Bandeiras from the Merck Healthcare KGaA Satellite Lab of the Structural Biology for Drug Discovery Unit for their help and great discussions. We thank Manuel Melo (ITQB NOVA) and Maria João Amorim (CRB) for important expert feedback. We thank the COVID task force initiative from ITQB NOVA and all its members.

## CONFLICT OF INTEREST

The authors declare no competing interest.

## Notes

### Competing Interest Statement

The authors have declared no competing interest.

