## Supplementary Information for "Dynamic ensembles of SARS-CoV-2 N-protein reveal head-to-head coiled-coil-driven oligomerization and phase separation"

#### FIGURE SI

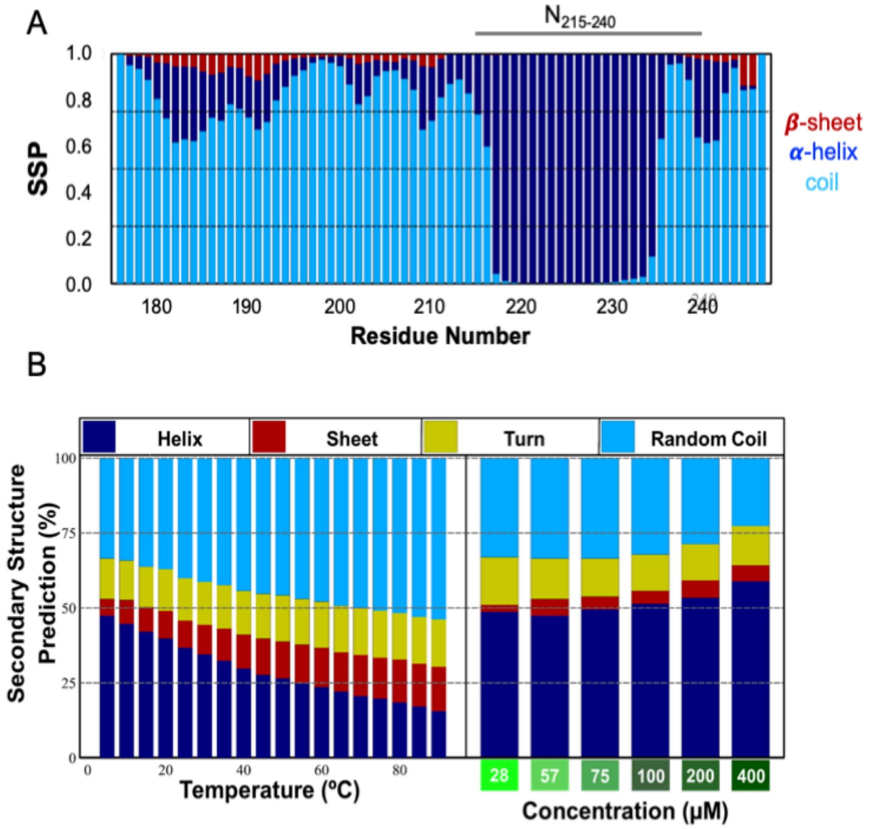


**Figure S1 - $N_{215\text{-}240}$ secondary structure predictions.** (**A**) Relative secondary structure propensity (SSP) predicted for IDL sequence, with β-sheet in red, α-helix in blue and random coil in green. (**B**) Prediction of $N_{215\text{-}240}$ secondary structure based on its far-UV CD spectra. Secondary structure content along the thermal denaturation (Left) and as a function of peptide concentration (Right). Helicity (dark blue) decreases with temperature and increases with peptide concentration.

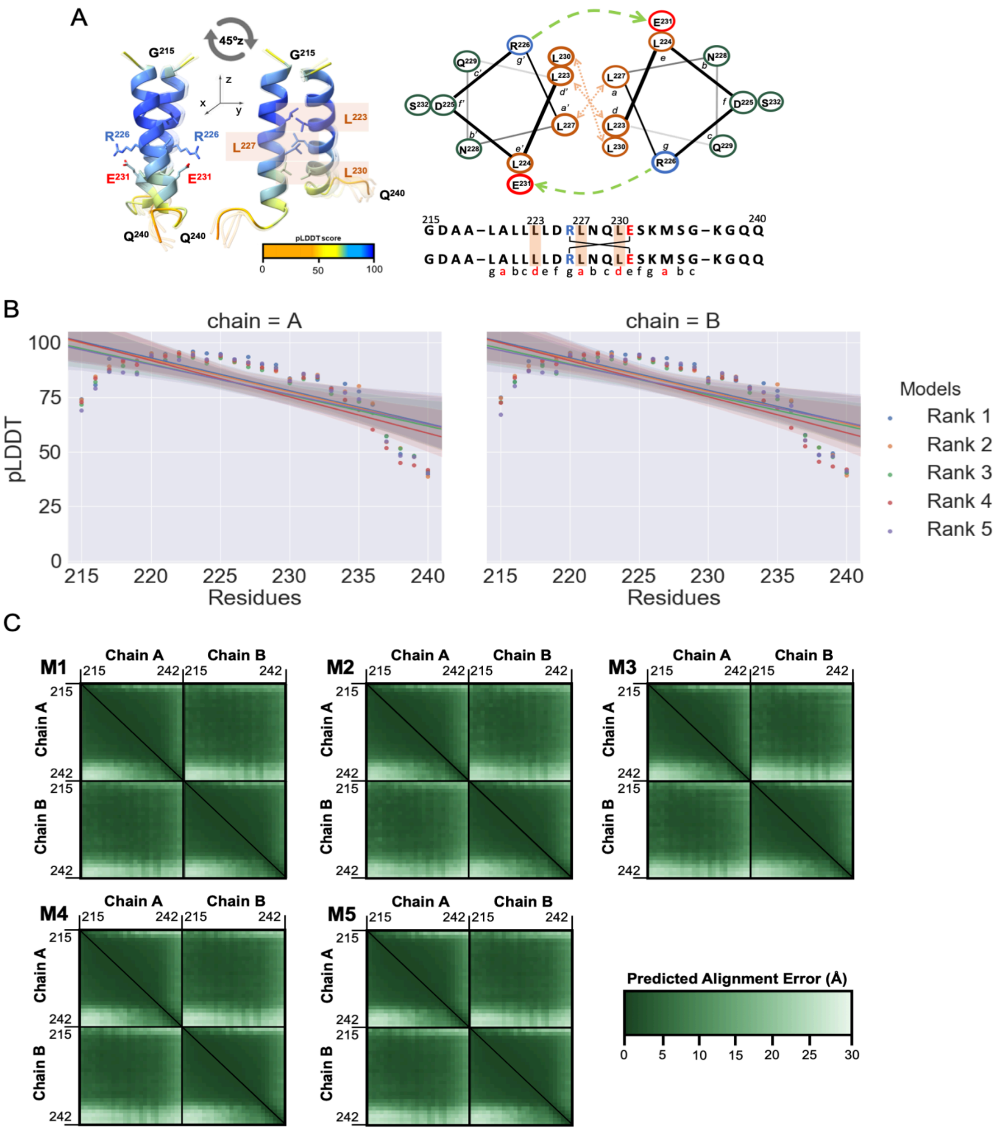


**Figure S2 - $N_{215\text{-}240}$ coiled-coil dimer prediction**. (**A**) Coiled-coil dimer predicted with AF-2 coloured according to pLDDT metric. We show an overlay of the five best models. The electrostatic pair R226-E231 and *knobs-into-holes* leucines 223, 227, and 230 are in stick representation. Helical wheel for the predicted homodimer labelled from *a* to *g*. Interface positions (*a*, *d*) are coloured orange, and solvent-exposed positions (*b*, *c* & *f*) are coloured green. Residues at *a* and *d* create a hydrophobic core, while *e* and *g* residues favour dimerisation through a stabilising salt bridge. The interfacial hydrophobic (*a, d*) and charged (*g, e*) residues, stabilizing a parallel orientation, are indicated in the primary sequence (bottom). (**B**) Profiles of pLDDT for chains A (left) and B (right) of the best-ranked five CC-Di models display a high mean pLDDT score (80.34±15.68). (**C**) Predicted Aligned Errors (PAE) on a scale from 0 to 30 Å in a green gradient. PAE is a metric of confidence in the relative position and orientation of the different chains of the model. All models display low inter-chain PAE values (green), indicative of well-defined relative inter-chain positioning within the predicted CC-dimer.

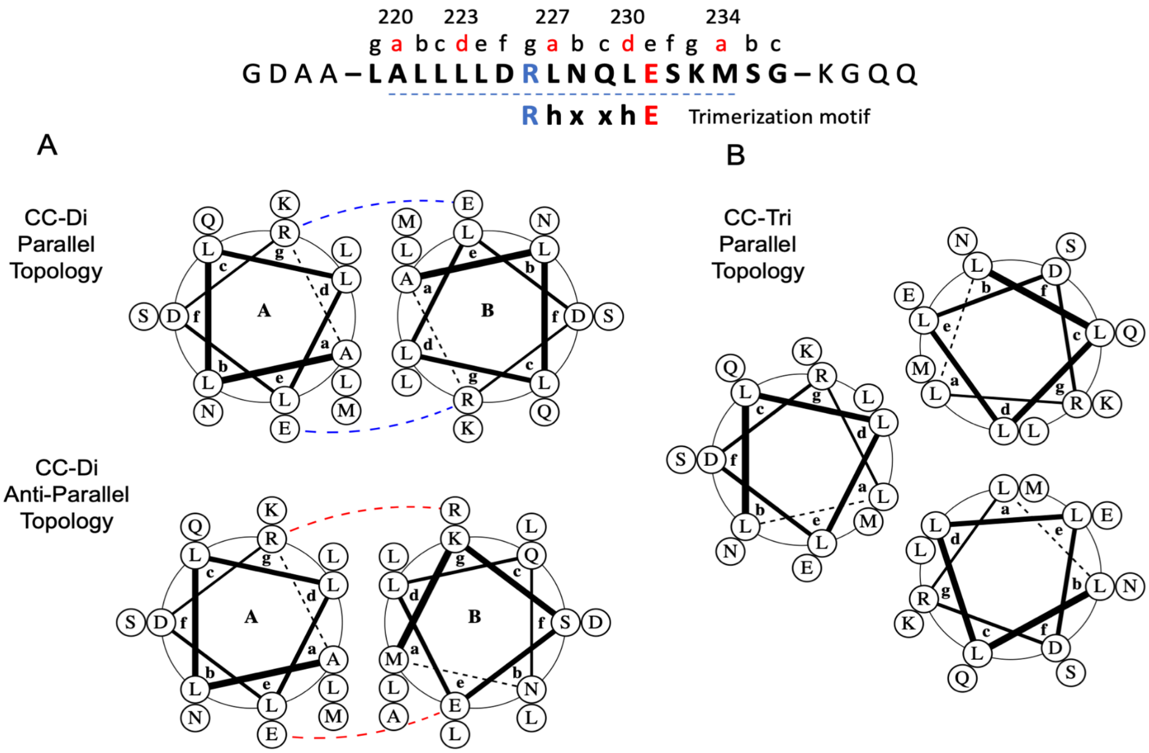


**Figure S3 - $N_{215-240}$ potential coiled-coil states.** (**A**) Helical wheel diagram for the $N_{215-240}$ CC-Di in parallel and anti-parallel topologies. Potential interfacial electrostatic bridges and repulsion interactions are indicated by blue and red dashed lines, respectively. In the parallel dimer, R226 and E231 form a salt bridge. (**B**) Helical wheel diagram for the parallel $N_{215-240}$ CC-Tri. The sequence $N_{215-240}$ (top) contains a trimerization motif, R226-h(a)xxh(d)E231 where h(a) represents hydrophobic residues such as Ile, Leu, Val, Met, and h(d) represents Leu, Ile or Val, with x being any amino acid residue. This motif specifies a three-stranded, parallel topology in short coiled coils.

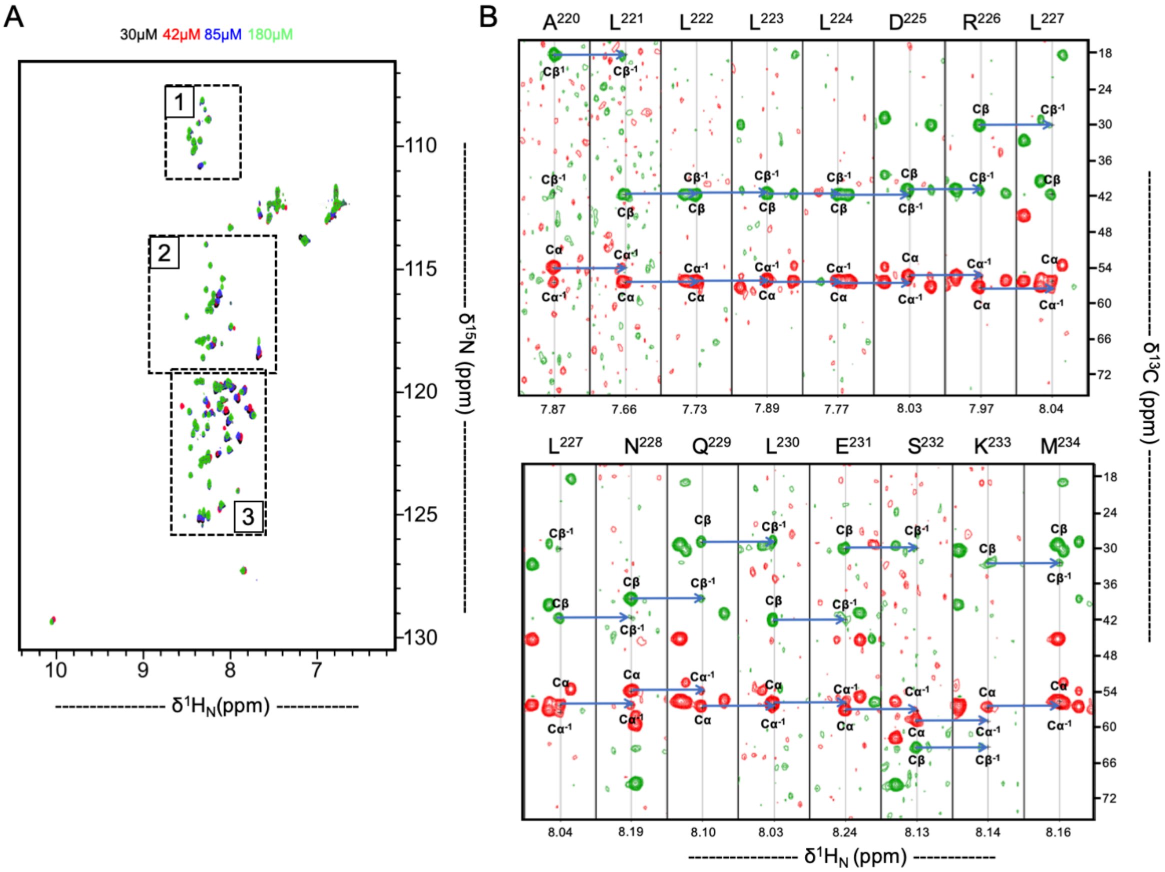


**Figure S4 - Interdomain linker resonances.** (**A**) [$^{15}$N-$^{1}$HN]-HSQC spectra of IDL ($N_{176\text{-}246}$) at different concentrations. Differential enhanced line-broadening and intensity drop are observed with concentration, consistent with structural heterogeneity and conformational exchange between monomers. Expanded views are shown in **Figure 3** for the regions within dashed boxes. All spectra display a narrow $^{1}$H chemical shift dispersion characteristic of disordered proteins with $^{1}$H amide backbone resonances clustering between 7.7 and 8.5 ppm. (**B**) Sequential correlations in the HNCACB experiment marked by horizontal blue arrows show ten different spin systems of the helical region found in the IDL. Notice the opposite sign for C$\alpha$ (red) and C$\beta$ (green) cross-peaks.

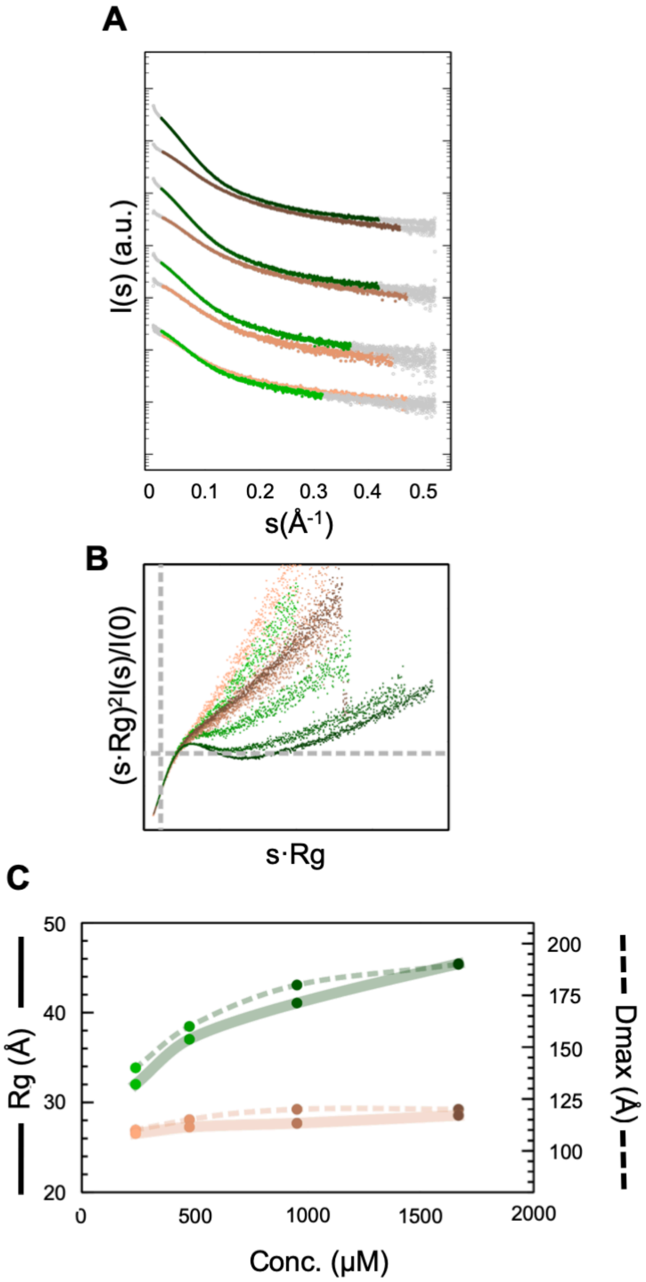


**Figure S5 - SAXS analysis of mutations' impact on IDL CC site.** (**A**) Logarithmic-scale representation of scattering intensity, *I(s)*, as a function of the momentum transfer, *s*, measured (grey circles) and used for IDL (green circles) and CC-null variant (brown) at different concentrations: 238.1, 476.2, 952.3, and 1666.7 μM. (**B**) Dimensionless Kratky representation of SAXS data measured for IDL (green) and CC-null (brown). The absence of a distinct peak maximum at $sRg = \sqrt{3}$ (dashed line) implies that the IDL remains conformationally heterogeneous even upon self-assembly. (**C**) *Rg* (solid lines) and $D_{max}$ (dashed lines evolution with concentration for IDL (green) and CC-null (brown).

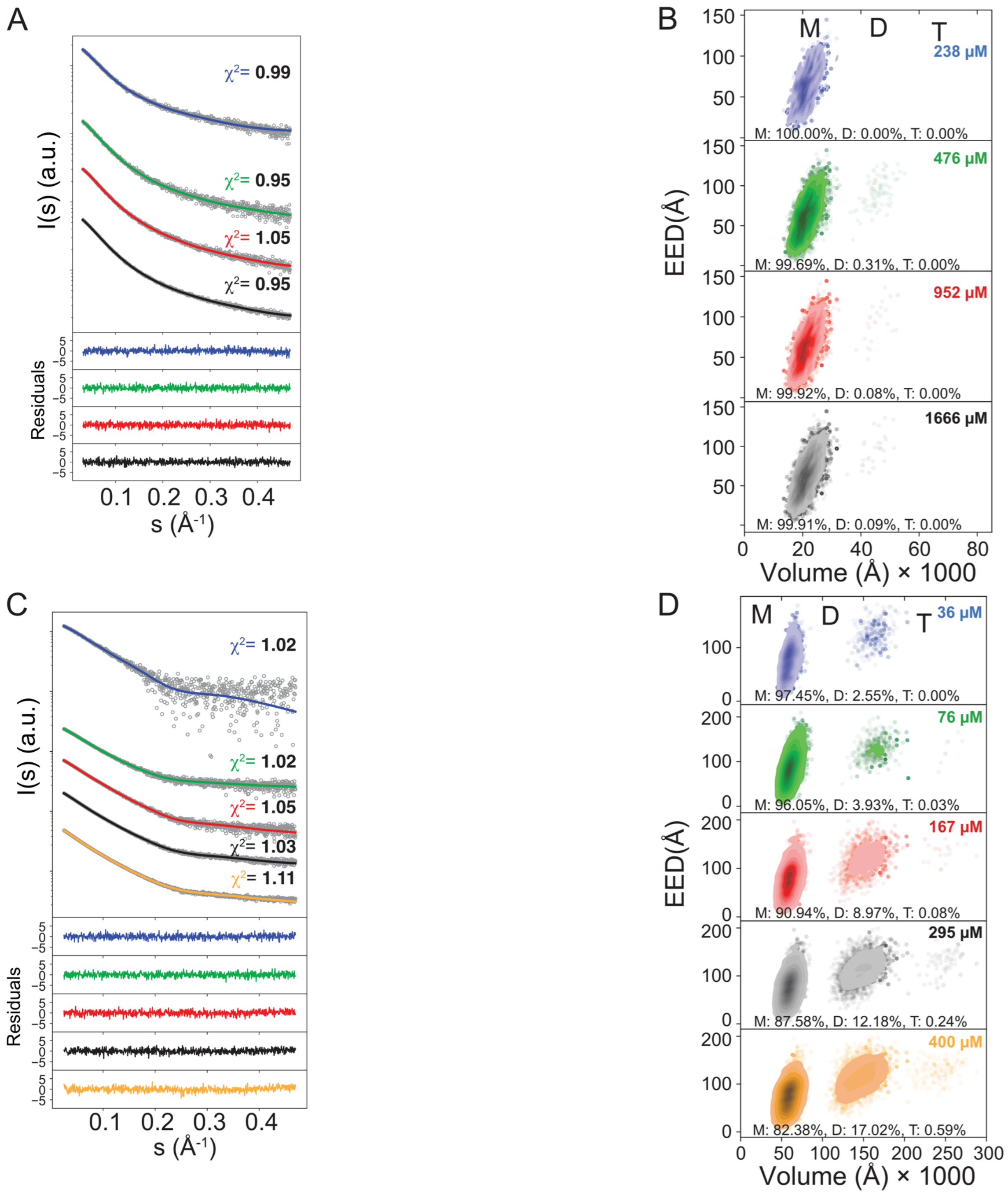


**Figure S6 - Impact of mutations on the monomer-to-trimer self-assembly of IDL and $N_{1\text{-}246}$ by targeting the CC site. (A, C)** Experimental SAXS profiles for $N_{176\text{-}246}$L3P and $N_{1\text{-}246}$L3P at different concentrations in grey and respective EOM fittings in solid lines. χ2 values are labelled next to each SAXS-curve. Point-by-point residuals of the fittings are at the bottom. (**B, D**) KDE contour plots for End-to-End distance (EED) and Volume, calculated from the EOM-selected sub-ensembles, are provided next to each SAXS curves panel and colored using the same color code. "M", "D", and “T” denote monomer, dimer and trimer species, respectively.

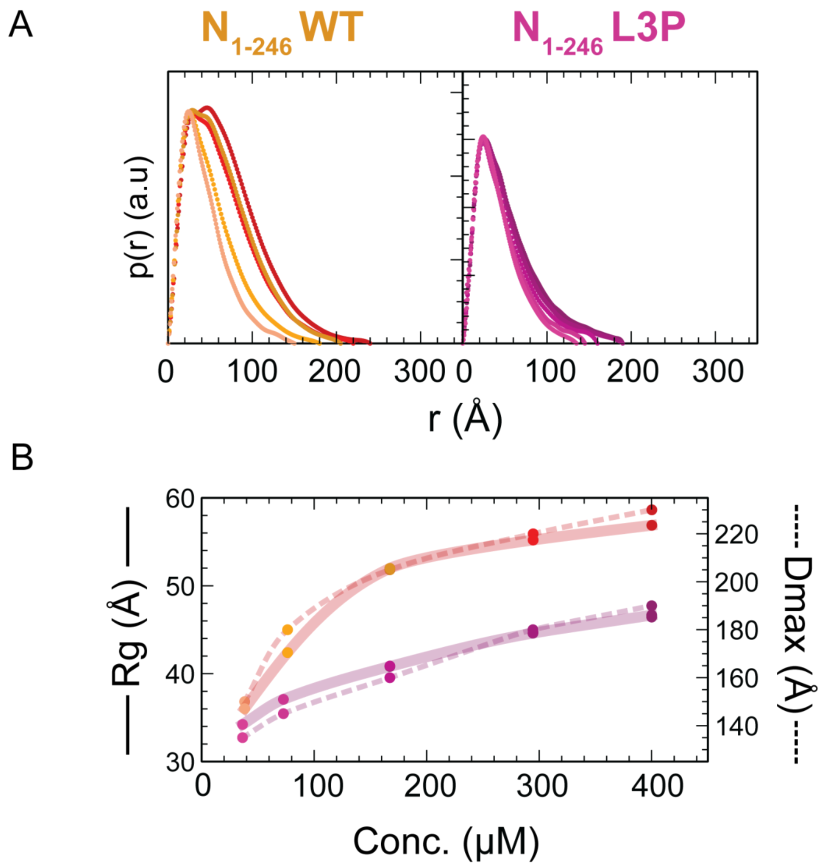


**Figure S7 - SAXS data analysis for $N_{1\text{-}246}$ WT and L3P protein constructs. (A)** Pair distance distributions, reporting all distances between any two scatterers within the SAXS profiles, retrieved from the SAXS profiles for $N_{1\text{-}246}$ WT (red gradient) and $N_{1\text{-}246}$ L3P (magenta gradient). Concentrations are indicated in **Supplementary Table 4** spanning from lowest (light colour) to highest (dark colour). **(B)** *Rg* (solid lines) and $D_{max}$ (dashed lines) at experimental concentrations for $N_{1\text{-}246}$ WT and $N_{1\text{-}246}$ L3P.

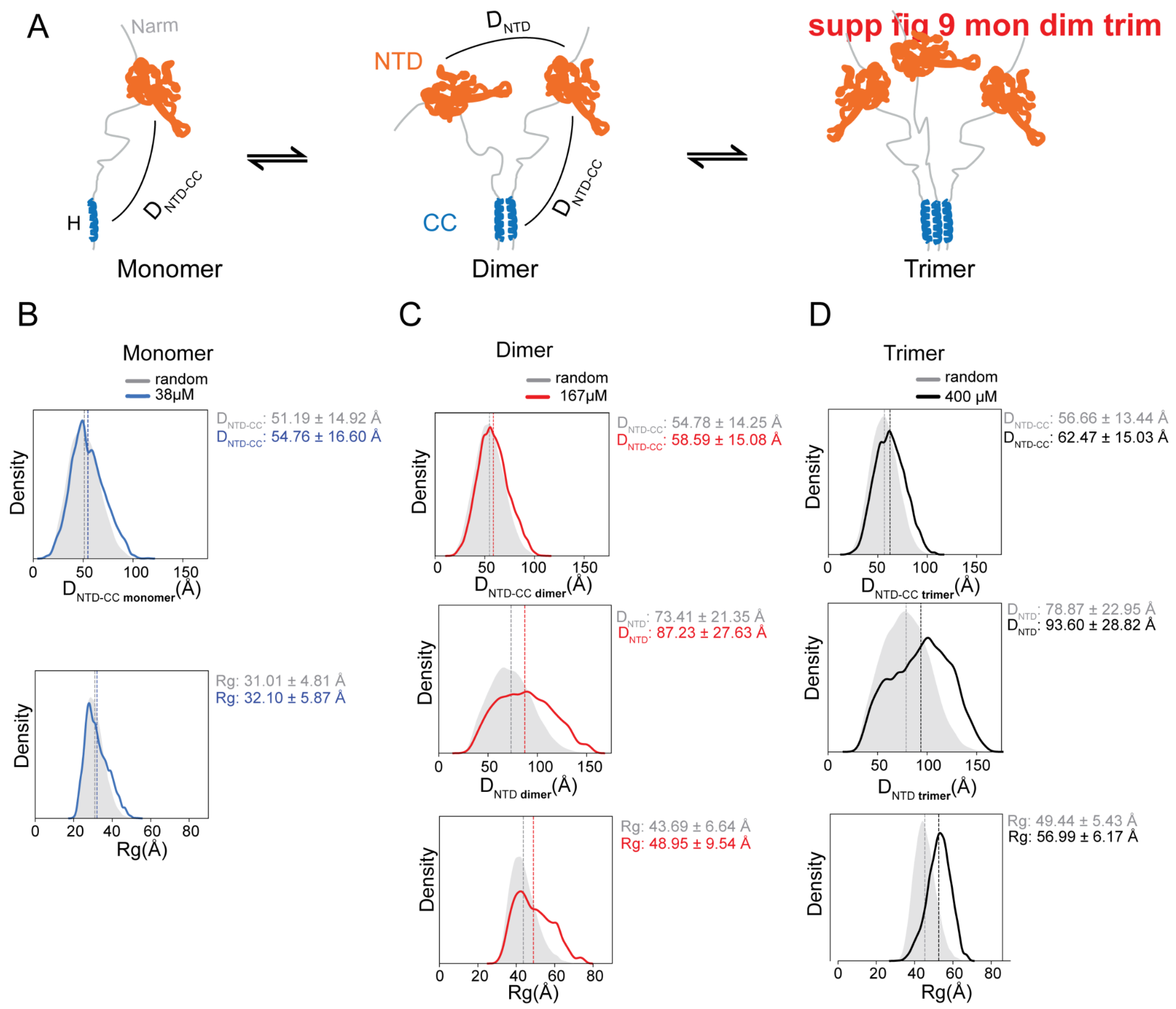


**Figure S8 - Structural distance distributions for the monomer-to-trimer assembly of $N_{1\text{-}246}$. (A)** Schema representing the distance distributions for the monomer-dimer equilibrium. $D_{NTD\text{-}CC}$ represents the distance between the CC domain and helix H, in blue, and the NTD domain, in orange. $D_{NTD}$ are the distances between NTD domains. The inter-domain linker and Narm tail are depicted in light grey. **(B)** Monomeric State Analysis: Kernel density estimation (KDE) plots depict the monomeric EOM-selected sub-ensembles for distances between CC and NTD domains ($D_{NTD\text{-}CC}$) and Rg at the indicated concentration. **(C,D)** KDE plots display the dimeric and trimeric EOM-selected sub-ensembles for the $D_{NTD\text{-}CC}$, $D_{NTD}$, and *Rg* distance distributions at the indicated concentrations. The filled grey area represents the KDE for the respective random ensembles. In dashed vertical lines is drawn the corresponding metric average. The average metric numerical values together with the standard deviation are also indicated inside plot borders.

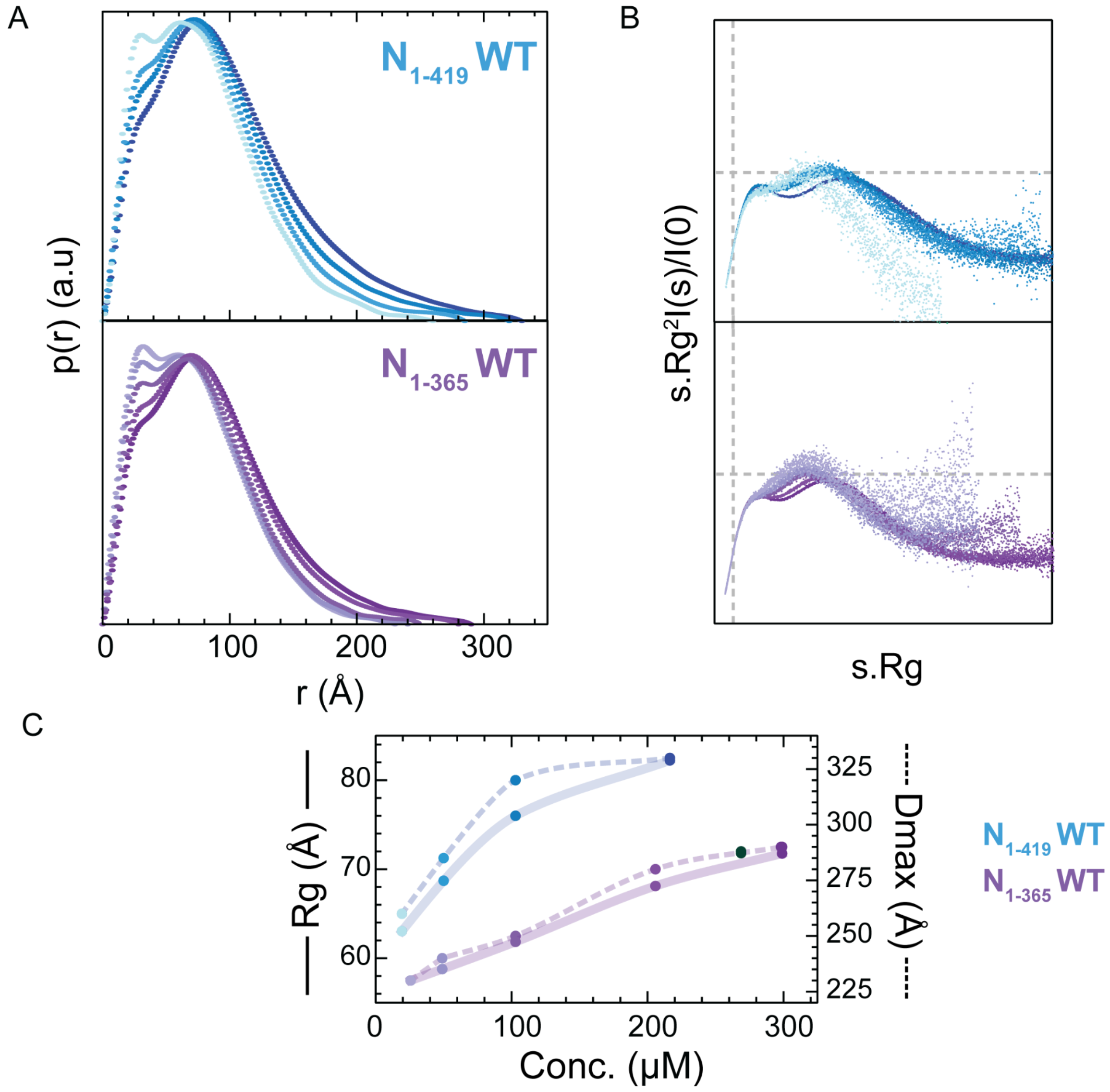


**Figure S9 - SAXS data analysis for $N_{1\text{-}419}$ and $N_{1\text{-}365}$ WT protein constructs. (A)** Pair distance distributions, reporting all distances between any two scatterers within the SAXS profiles, retrieved from the SAXS profiles for $N_{1\text{-}419}$ (blue gradient) and $N_{1\text{-}365}$ (purple gradient). Concentrations are indicated in **Supplementary Table 4** spanning from lowest (light colour) to highest (dark colour). **(B)** Dimensionless Kratky representation of SAXS data measured for $N_{1\text{-}419}$ and $N_{1\text{-}365}$. The absence of a distinct peak maximum at $sRg = \sqrt{3}$ (dashed line) implies that all proteins remain conformationally heterogeneous even upon self-assembly. The colour code is the same as in panel (A). (**C**) *Rg* (solid lines) and $D_{max}$ (dashed lines evolution with concentration for $N_{1\text{-}419}$ (in blue) and $N_{1\text{-}365}$ (in purple).

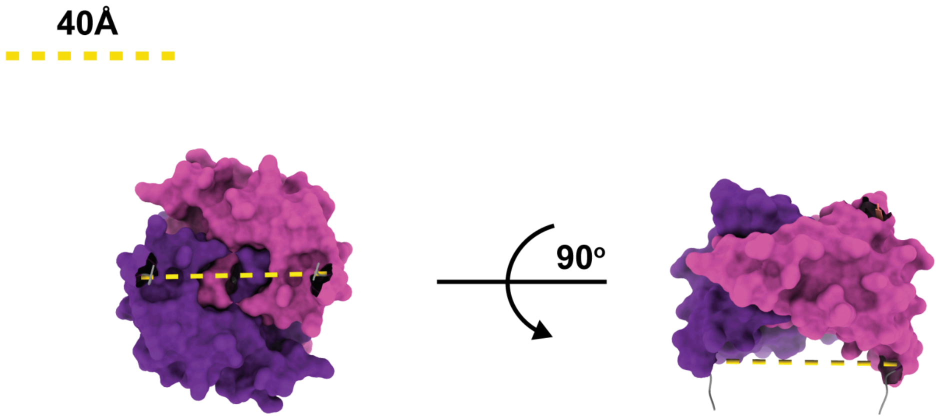


**Figure S10 - Distance between the N-terminal of individual monomers for the CTD domain dimer**. A yellow dashed line represents the Cα-Cα distance between the N-terminal residue S250 of the CTD anti-parallel swapped dimer. The CTD dimer is depicted in surface representation with light and dark purple chains. The right panel represents the CTD with a 90º rotation.

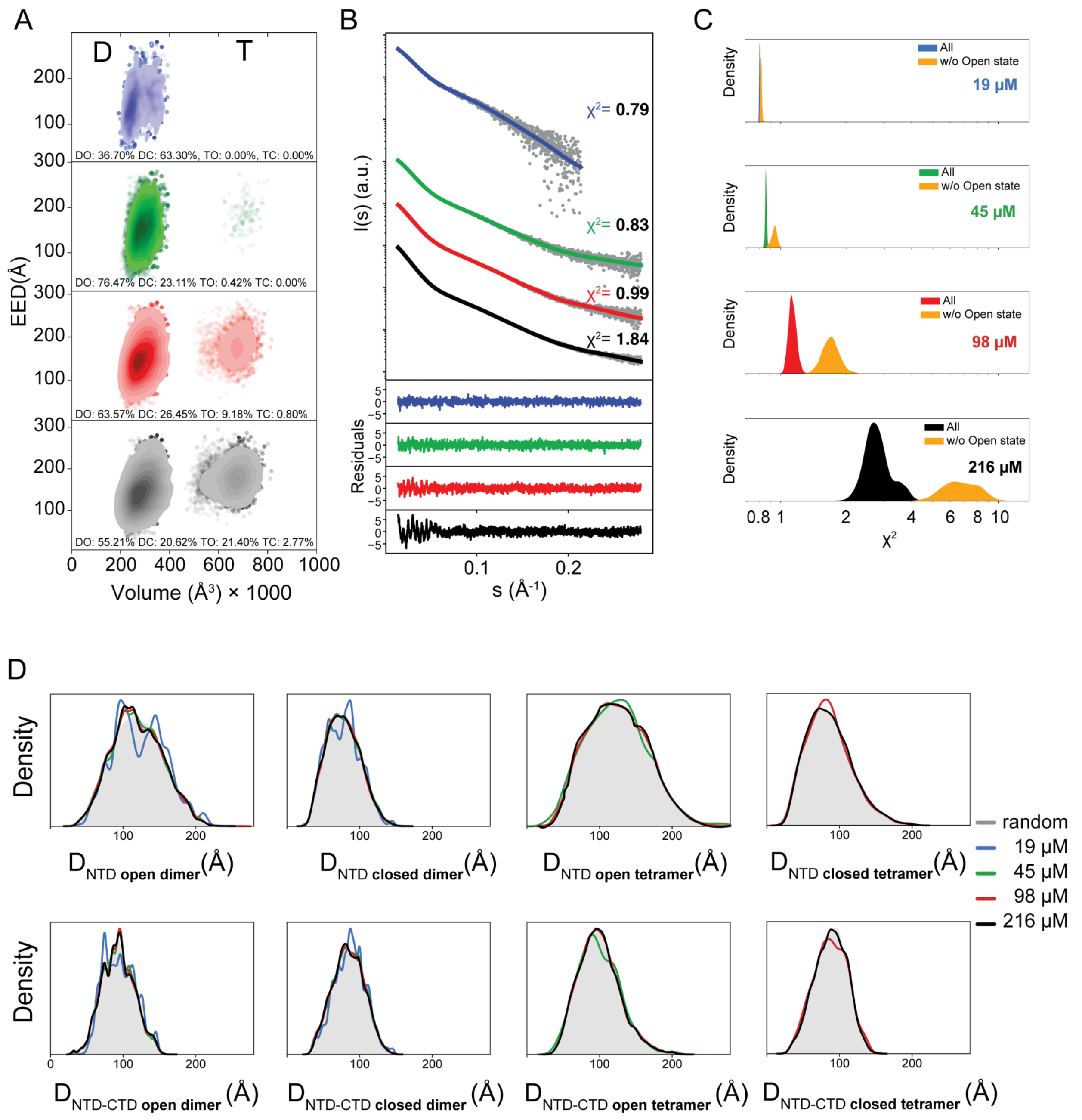


**Figure S11 - SAXS data analysis for dimer-tetramer equilibrium of** $N_{1\text{-}419}$**.**

(**A**) KDE contour plots showing the End-to-End Distance (EED) and Volume distributions, calculated for the EOM-selected sub-ensembles at concentrations of 19 (blue), 45 (green), 98 (red), and 216 µM (black). The dimer (D) and tetramer (T) species are represented, with DO, DC, TO, and TC indicating dimer open, dimer closed, tetramer open, and tetramer closed states, respectively. (**B**) KDE plots of χ2-values from 2000 EOM trial runs for each concentration. The light yellow distribution represents the χ2 values excluding the open dimeric and tetrameric states, while the distributions including all four states

(DO, DC, TO, TC) are shown in the same color scheme as panel A. (**C**) Experimental SAXS profiles for $N_{1\text{-}419}$ at different concentrations (grey dots), overlaid with the corresponding EOM fits (solid lines) using the colour code as in panel A. The $\chi 2$ values for the fits are indicated next to each SAXS curve, with the point-by-point residuals displayed beneath the profiles. **(D)** KDE plots of the $D_{NTD}$ and $D_{NTD\text{-}CTD}$ distributions for EOM-selected open and closed dimeric and tetrameric states. These plots correspond to the same concentrations and color codes as in panel C. The filled grey areas represent the KDEs for the respective random ensembles, serving as a reference for structural selection.

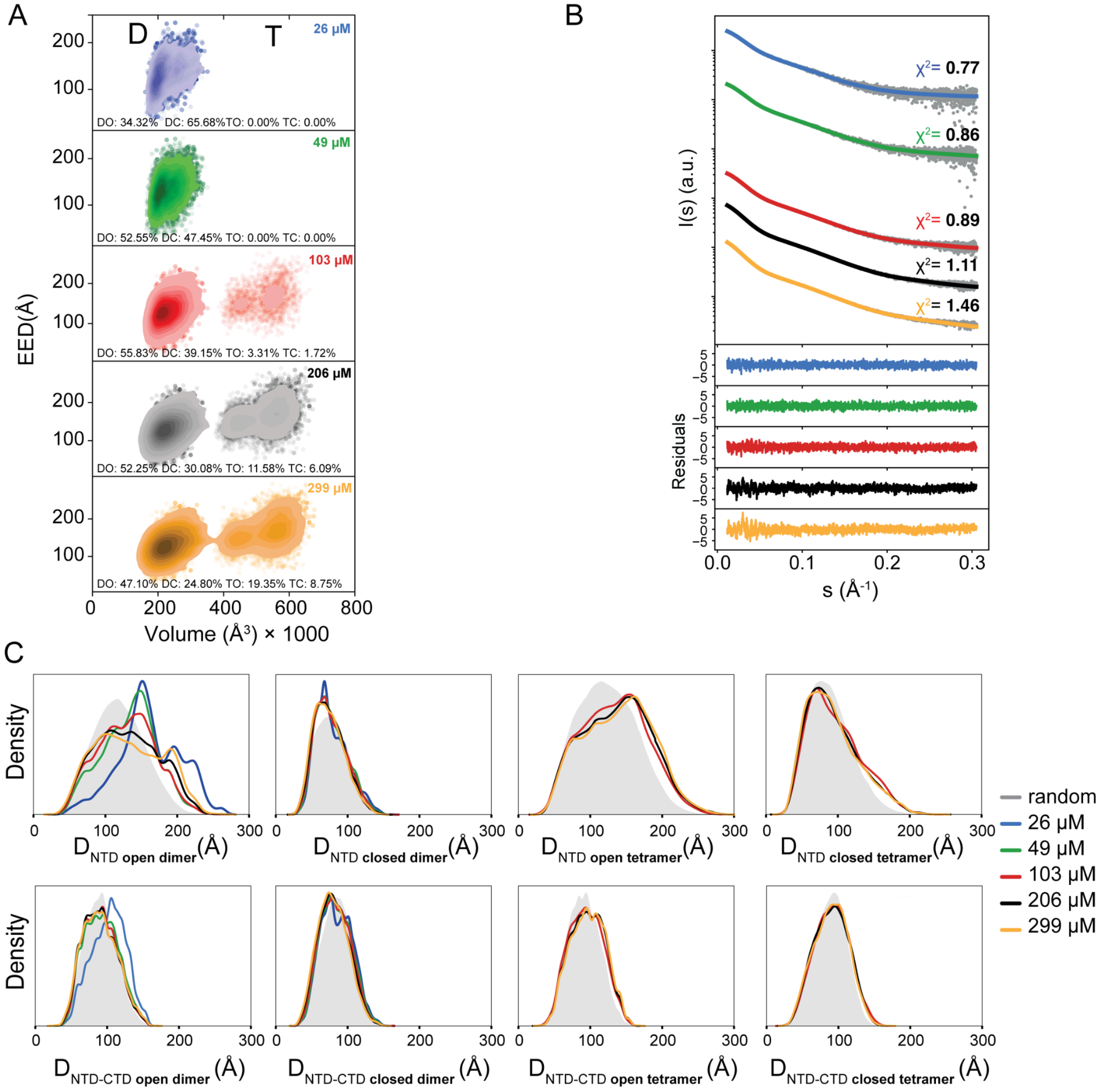


**Figure S12 - SAXS data analysis for dimer-tetramer equilibrium of** $N_{1\text{-}365}$**.** (**A**) KDE contour plots for the End-to-End Distance (EED) and Volume, calculated for the EOM-selected sub-ensembles at concentrations of 26, 49, 103, 206 and 299 μM, shown in blue, green, red, black, and yellow, respectively. D, and T denote dimer and tetramer species, respectively. DO, DC, TO and TC denote dimer open, dimer closed, tetramer open and tetramer closed, respectively. **(B)** Experimental SAXS profiles for $N_{1\text{-}365}$ at different concentrations in grey and respective EOM fittings in solid lines. χ2 values are labelled next to each SAXS-curve. Point-by-point residuals of the fittings are at the bottom. Color code is the same as in panel A. **(C)** KDE plots of the DNTD and DNTD-CTD distance distributions for EOM-selected open and closed dimeric and tetrameric states at varying concentrations. The filled grey areas represent the KDEs for the respective random ensembles.

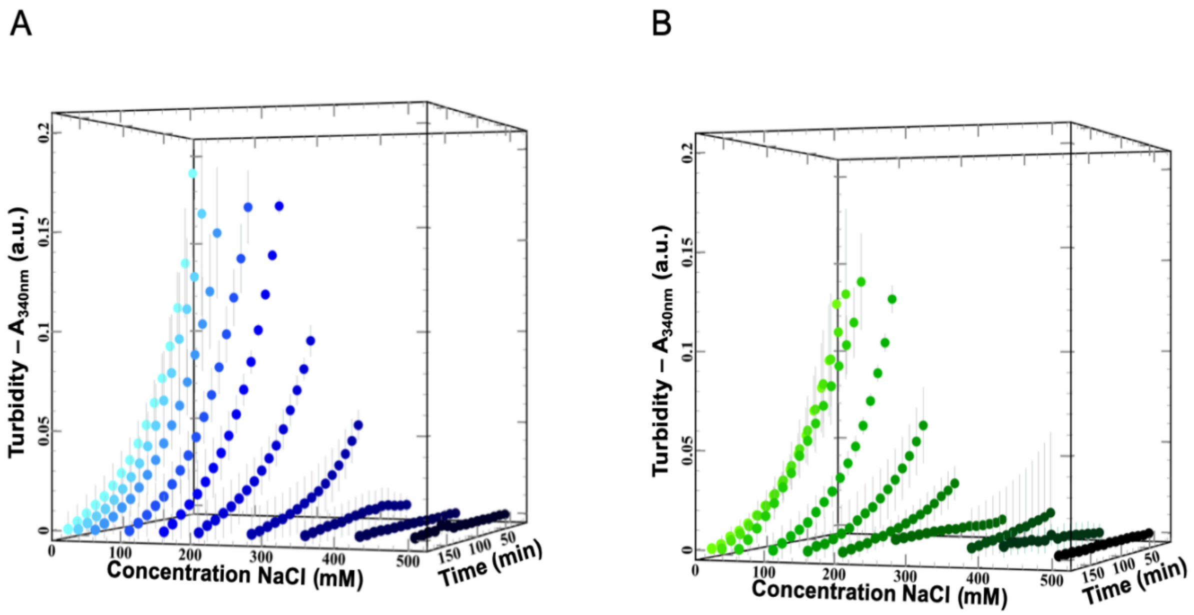


**Figure S13 - Phase separation over time as monitored by turbidity of $N_{1\text{-}419}$ and $N_{1\text{-}419}$L3P.** (**A**) 3D-representation of turbidity measurements of $N_{1\text{-}419}$ at different salt concentrations over time. (**B**) 3D-representation turbidity measurements at 340 nm of $N_{1\text{-}419}$L3P at different salt concentrations over time. All turbidity traces are mean ± SD of experiments in triplicate. Please note: 2D projections (contour plots as a function of time/[NaCl]) of these data are shown in **Figure 7**.

### SUPPLEMENTARY TABLES

**Supplementary Table 1.** Primers used in this study.

| **Primer** | **Sequence (from 5' to 3')**<br>Lowercase = non-priming overhangs<br>Underlined = HRV 3C-Protease cleavage site<br>Bold= StrepTag + Stop codon<br>Uppercase= Gene specific |
|---|---|
| **NP_1_for** | tca gca agg gct gag gCT CGA GGT ACT CTT TCA AGG ACC GAT GTC AGA TAA TGG GCC TCA AAA CCA G |
| **NP_419_rev** | tca gcg gaa gct gag g**CT ACT TCT CGA ATT GAG GGT GAC TCC A**AG CCT GCG TAG AGT CCG CAG A |
| **NP_176_for** | tca gca agg gct gag gCT CGA GGT ACT CTT TCA AGG ACC GTC GCG TGG TGG CAG CCA G |
| **NP_246_rev** | tca gcg gaa gct gag g**CT ACT TCT CGA ATT GAG GGT GAC TCC A**AA CGG TCT GGC CCT GCT G |
| **NP_365_rev** | tca gcg gaa gct gag g**CT ACT TCT CGA ATT GAG GGT GAC TCC A**CG TTA CTT CCA TGC CGA TGC G |

**Supplementary Table 2.** Protein constructs expressed and puried in this study.

| **Construct** | **Protein sequence**[a]<br>Bold= linker sequence / Red = Mutations / Green = Green fluorescence protein |
|---|---|
| Sars-CoV-2 Nucleocapsid protein variants | |
| **$N_{176-276}$** | *GP*-**176SRGGSQASSRSSSRSRNSSRNSTPGSSRGTSPARMAGNGGDAALALLLLDRLNQLESKMSGKGQQQQGQTV246**-*WSHPQFEK* |
| **$N_{176-276}$L3P** | *GP*-**176SRGGSQASSRSSSRSRNSSRNSTPGSSRGTSPARMAGNGGDAALALLPLDRPNQPESKMSGKGQQQQGQTV246**-*WSHPQFEK* |
| **$N_{1-246}$** | *GP*-**1**MSDNGPQNQRNAPRITFGGPSDSTGSNQNGERSGARSKQRRPQGLPNNTASWFTALTQHGKEDLKFPRGQGVPINTNSSPDDQIGYYRRATRRIRGGDGKMKDLSPRWYFYYLGTGPEAGLPYGANKDGIIWVATEGALNTPKDHIGTRNPANNAAIVLQLPQGTTLPKGFYAEG**SRGGSQASSRSSSRSRNSSRNSTPGSSRGTSPARMAGNGGDAALALLLLDRLNQLESKMSGKGQQQQGQTV246**-*WSHPQFEK* |
| **$N_{1-246}$L3P** | *GP*-**1**MSDNGPQNQRNAPRITFGGPSDSTGSNQNGERSGARSKQRRPQGLPNNTASWFTALTQHGKEDLKFPRGQGVPINTNSSPDDQIGYYRRATRRIRGGDGKMKDLSPRWYFYYLGTGPEAGLPYGANKDGIIWVATEGALNTPKDHIGTRNPANNAAIVLQL |

| | |
|---|---|
| | PQGTTLPKGFYAEG**SRGGSQASSRSSSRSRNSSRNSTPGSSRGTSPARMAGNG**<br>**GDAALALLPLDRPNQPESKMSGKGQQQQGQTV246**-*WSHPQFEK* |
| **N$_{1-365}$** | *GP*-**1**MSDNGPQNQRNAPRITFGGPSDSTGSNQNGERSGARSKQRRPQGLPNNTA<br>SWFTALTQHGKEDLKFPRGQGVPINTNSSPDDQIGYYRRATRRIRGGDGKMKDLS<br>PRWYFYYLGTGPEAGLPYGANKDGIIWVATEGALNTPKDHIGTRNPANNAAIVLQL<br>PQGTTLPKGFYAEG**SRGGSQASSRSSSRSRNSSRNSTPGSSRGTSPARMAGNG**<br>**GDAALALLLLDRLNQLESKMSGKGQQQQGQTV**TKKSAAEASKKPRQKRTATKAY<br>NVTQAFGRRGPEQTQGNFGDQELIRQGTDYKHWPQIAQFAPSASAFFGMSRIGM<br>EVTPSGTWLTYTGAIKLDDKDPNFKDQVILLNKHIDAYKTFPP**365**-*WSHPQFEK* |
| **N$_{1-419}$** | *GP*-**1**MSDNGPQNQRNAPRITFGGPSDSTGSNQNGERSGARSKQRRPQGLPNNTA<br>SWFTALTQHGKEDLKFPRGQGVPINTNSSPDDQIGYYRRATRRIRGGDGKMKDLS<br>PRWYFYYLGTGPEAGLPYGANKDGIIWVATEGALNTPKDHIGTRNPANNAAIVLQL<br>PQGTTLPKGFYAEG**SRGGSQASSRSSSRSRNSSRNSTPGSSRGTSPARMAGNG**<br>**GDAALALLLLDRLNQLESKMSGKGQQQQGQTV**TKKSAAEASKKPRQKRTATKAY<br>NVTQAFGRRGPEQTQGNFGDQELIRQGTDYKHWPQIAQFAPSASAFFGMSRIGM<br>EVTPSGTWLTYTGAIKLDDKDPNFKDQVILLNKHIDAYKTFPPTEPKKDKKKKADET<br>QALPQRQKKQQTVTLLPAADLDDFSKQLQQSMSSADSTQA**419-***WSHPQFEK* |
| **GFP-N$_{1-419}$** | *MG*VSKGEELFTGVVPILVELDGDVNGHKFSVSGEGEGDATYGKLTLKFICTTGKLP<br>*VPWPTLVTTLTYGVQCFARYPDHMKQHDFFKSAMPEGYVQERTIFFKDDGNYKTR*<br>*AEVKFEGDTLVNRIELKGIDFKEDGNILGHKLEYNYNSHKVYITADKQKNGIKVNFKT*<br>*RHNIEDGSVQLADHYQQNTPIGDGPVLLPDNHYLSTQSALSKDPNEKRDHMVLLE*<br>*FVTAAGITLGMDELYKSSGPSGSSHHHHHHSSGPQQGLRLEVLFQGP*-**1**MSDNGP<br>QNQRNAPRITFGGPSDSTGSNQNGERSGARSKQRRPQGLPNNTASWFTALTQHG<br>KEDLKFPRGQGVPINTNSSPDDQIGYYRRATRRIRGGDGKMKDLSPRWYFYYLGT<br>GPEAGLPYGANKDGIIWVATEGALNTPKDHIGTRNPANNAAIVLQLPQGTTLPKGFY<br>AEG**SRGGSQASSRSSSRSRNSSRNSTPGSSRGTSPARMAGNGGDAALALLLLD**<br>**RLNQLESKMSGKGQQQQGQTV**TKKSAAEASKKPRQKRTATKAYNVTQAFGRRG<br>PEQTQGNFGDQELIRQGTDYKHWPQIAQFAPSASAFFGMSRIGMEVTPSGTWLTY<br>TGAIKLDDKDPNFKDQVILLNKHIDAYKTFPPTEPKKDKKKKADETQALPQRQKKQ<br>QTVTLLPAADLDDFSKQLQQSMSSADSTQA**419-***WSHPQFEK* |
| **GFP-N$_{1-419}$L3P** | *MGVSKGEELFTGVVPILVELDGDVNGHKFSVSGEGEGDATYGKLTLKFICTTGKLP*<br>*VPWPTLVTTLTYGVQCFARYPDHMKQHDFFKSAMPEGYVQERTIFFKDDGNYKTR*<br>*AEVKFEGDTLVNRIELKGIDFKEDGNILGHKLEYNYNSHKVYITADKQKNGIKVNFKT*<br>*RHNIEDGSVQLADHYQQNTPIGDGPVLLPDNHYLSTQSALSKDPNEKRDHMVLLE*<br>*FVTAAGITLGMDELYKSSGPSGSSHHHHHHSSGPQQGLRLEVLFQGP*-**1**MSDNGP<br>QNQRNAPRITFGGPSDSTGSNQNGERSGARSKQRRPQGLPNNTASWFTALTQHG<br>KEDLKFPRGQGVPINTNSSPDDQIGYYRRATRRIRGGDGKMKDLSPRWYFYYLGT<br>GPEAGLPYGANKDGIIWVATEGALNTPKDHIGTRNPANNAAIVLQLPQGTTLPKGFY<br>AEG**SRGGSQASSRSSSRSRNSSRNSTPGSSRGTSPARMAGNGGDAALALLPLD**<br>**RPNQPESKMSGKGQQQQGQTV**TKKSAAEASKKPRQKRTATKAYNVTQAFGRRG<br>PEQTQGNFGDQELIRQGTDYKHWPQIAQFAPSASAFFGMSRIGMEVTPSGTWLTY<br>TGAIKLDDKDPNFKDQVILLNKHIDAYKTFPPTEPKKDKKKKADETQALPQRQKKQ<br>QTVTLLPAADLDDFSKQLQQSMSSADSTQA**419-***WSHPQFEK* |

[a]The first and last residues are in bold and extra residues due to cloning are in italics.

**Supplementary Table 3. Bio-SAXS Beamline and acquisition parameters for N-Protein constructions.**

| Construction | $N_{1-419}$ | $N_{1-365}$ | $N_{1-246}$ | | $N_{176-236}$ | |
|---|---|---|---|---|---|---|
| | Wild-Type | Wild-Type | Wild-Type | $L^{223,227,230}P$ | Wild-Type | $L^{223,227,230}P$ |
| Experimental Session | MX25270-8 | MX25270-8 | MX25270-8 | MX-2450 | MX-2450 | MX-2450 |
| Beamline | B21 – DSL | | | BM29 – ERF | | |
| Wavelength (Å) | 0.954 | | | 0.999 | | |
| Sample-to-detector distance (mm) | 3688.3 | | | 2813.0 | | |
| *s* range ($Å^{-1}$) | 0.0045-0.34 | | | 0.0053-0.52 | | |
| HPLC system | Agilent 1200 HPLC System | | | Shimazu HPLC System | | |
| SEC column | Shodex KW403-4F | | Shodex KW402.5-4F | Agilent BIO-SEC-3 4.6/300 | | |
| Detector | EigerX 4M (Dectris) | | | Pilatus 1M | | |
| Temperature (K) | 288.15 | | | | | |

Software

| | | | |
|---|---|---|---|
| SEC-SAXS data integration | ScÅtter3 | *P(r)* | GNOM 5.0 |

**Supplementary Table 4. SAXS-derived parameters from N-Protein samples.**

| | Concentration (mg/mL // μM) | SASBDB Codes | *s* range (Å$^{-1}$) | $R_g$ (Å) [from *P(r)*] | $R_g$ (Å) [from Guinier] | $D_{max}$ (Å) |
|---|---|---|---|---|---|---|
| **$N_{1\text{-}419}$ Wild-Type** | 0.9 // 19.3 | SASDTX5 | 0.0109-0.318 | 62.66 | 63.03±0.82 | 260±5 |
| | 2.1 // 50.0 | SASDTY5 | 0.0109-0.279 | 68.70 | 69.18±0.38 | 285±5 |
| | 4.8 // 102.9 | SASDTZ5 | 0.0109-0.299 | 75.42 | 76.00±0.30 | 320±5 |
| | 10.1 // 216.4 | SASDT26 | 0.0109-0.318 | 81.62 | 82.22±0.16 | 330±5 |
| **$N_{1\text{-}365}$ Wild-Type** | 1.0 // 25.7 | SASDTE5 | 0.0109-0.266 | 57.18 | 57.49±0.37 | 230±5 |
| | 2.0 // 49.0 | SASDTF5 | 0.0109-0.266 | 58.47 | 58.78±0.31 | 240±5 |
| | 4.2 // 102.9 | SASDTG5 | 0.0109-0.292 | 61.49 | 61.84±0.17 | 250±5 |
| | 8.4 // 205.8 | SASDTH5 | 0.0109-0.318 | 67.66 | 68.11±0.15 | 280±5 |
| | 12.2 // 299.0 | SASDTJ5 | 0.0109-0.331 | 71.29 | 71.76±0.10 | 290±5 |
| **$N_{1\text{-}246}$ Wild-Type** | 1.1 // 38.2 | SASDSR9 | 0.015-0.266 | 35.71 | 36.02±0.22 | 150±5 |
| | 2.1 // 76.4 | SASDSS9 | 0.015-0.266 | 42.05 | 42.40±0.17 | 180±5 |
| | 4.6 // 167.3 | SASDST9 | 0.015-0.305 | 52.01 | 52.42±0.13 | 220±5 |
| | 11.0 // 400.1 | SASDSU9 | 0.015-0.331 | 56.49 | 56.88±0.12 | 240±5 |
| **$N_{1\text{-}246}$L3P** | 1.0 // 36.4 | SASDSV9 | 0.0125-0.315 | 34.05 | 34.23±0.34 | 135±5 |
| | 2.0 // 72.7 | SASDSW9 | 0.0125-0.366 | 36.72 | 37.08±0.24 | 145±5 |

| | | | | | | |
|---|---|---|---|---|---|---|
| | 4.6 // 167.3 | SASDVU6 | 0.0125-0.418 | 40.44 | 40.87±0.16 | 160±5 |
| | 8.1 // 294.5 | SASDSX9 | 0.0125-0.469 | 44.14 | 44.61±0.18 | 190±5 |
| | 11.0 // 400.0 | SASDSY9 | 0.0125-0.469 | 45.94 | 46.44±0.15 | 190±5 |
| **$N_{176-246}$ Wild-Type** | 2.0 // 238.1 | SASDSZ9 | 0.0203-0.315 | 31.69 | 30.02±0.62 | 140±5 |
| | 4.0 // 476.2 | SASDT22 | 0.0203-0.366 | 36.60 | 37.03±0.54 | 160±5 |
| | 8.0 // 952.2 | SASDT32 | 0.0203-0.418 | 40.66 | 41.07±0.41 | 180±5 |
| | 14.0//1666.7 | SASDVS6 | 0.0233-0.418 | 44.98 | 45.45±0.24 | 190±5 |
| **$N_{176-246}$L3P** | 2.0 // 238.1 | SASDT42 | 0.0229-0.469 | 26.31 | 25.56±0.38 | 110±5 |
| | 4.0 // 476.2 | SASDT52 | 0.0229-0.444 | 27.00 | 27.25±0.48 | 115±5 |
| | 8.0 // 952.2 | SASDT62 | 0.0229-0.469 | 27.44 | 27.68±0.31 | 120±5 |
| | 14.0 // 1666.7 | SASDVT6 | 0.0229-0.456 | 28.24 | 28.55±0.21 | 120±5 |

**Supplementary Table 5. Protein Ensemble Database (PED) Codes for Each Oligomeric State[a]**

| Protein construct | Oligomeric State | PED Identifier |
|---|---|---|
| $N_{1-419}$ | 2-mer Open | |
| | 2-mer Closed | |
| | 4-mer Open | |
| | 4-mer Closed | |
| $N_{1-365}$ | 2-mer Open | |
| | 2-mer Closed | |
| | 4-mer Open | |
| | 4-mer Closed | |
| $N_{1-246}$ | Monomer | |
| | Dimer | |
| | Trimer | |
| $N_{176-246}$ | Monomer | |
| | Dimer | |
| | Trimer | |

[a] Each entry contains a sub-ensemble per concentration.

**Supplementary Table 6.** Turbidity over time parameters

| **$N_{1\text{-}419}$ Variant** | **Wild-type** | | **L3P** | |
|---|---|---|---|---|
| [NaCl] (mM) | 25 | 150 | 25 | 150 |
| $Turb(t = 0)$ | 0.256 ± 0.062 | 0.256 ± 0.011 | 0.199 ± 0.041 | 0.087 ± 0.022 |
| $Turb(t = \infty)$ | 0.273 ± 0.012 | 0.287 ± 0.014 | 0.271 ± 0.018 | 0.289 ± 0.022 |
| $k$ | 0.018 ± 0.005 | 0.015 ± 0.002 | 0.014 ± 0.006 | 0.014 ± 0.001 |
